# Phenologically-informed species distribution models (SDMs) forecast less species loss and turnover versus standard SDMs

**DOI:** 10.1101/2023.07.05.547862

**Authors:** Shijia Peng, Tadeo H. Ramirez-Parada, Susan Mazer, Sydne Record, Isaac Park, Aaron M. Ellison, Charles C. Davis

## Abstract

Species distribution models (SDMs) have been central for documenting the relationship between species’ geographic ranges and environmental conditions for more than two decades. However, the vast majority of SDMs rarely consider functional traits, such as phenology, which strongly affect species’ demography and fitness. Using >120,000 herbarium specimens representing 360 plant species across the eastern United States, we developed a novel “phenology-informed” SDM that integrates dynamic phenological responses to changing climates. Compared to standard SDMs based only on abiotic variables, our phenology-informed SDMs forecast significantly lower species habitat loss and less species turnover within communities under climate change. These results suggest that phenotypic plasticity or local adaptation in phenology may help many species adjust their ecological niches and persist in their habitats during periods of rapid environmental change. By modeling historical data that link phenology, climate, and species distributions, our findings reveal how species’ reproductive phenology mediates their geographic distributions along environmental gradients and affects regional biodiversity patterns in the face of future climate change. More importantly, our newly developed model also circumvents the need for mechanistic models that explicitly link traits to occurrences for each species, thus better facilitating the deployment of trait-based SDMs across unprecedented spatial and taxonomic scales.

## Main

The geographic distribution of species, especially plants and other sessile organisms, is strongly influenced by climate^1–4^. As global anthropogenic change intensifies, a better understanding of the climatic, environmental, and biotic factors that shape species distributions at large geographic, temporal, and taxonomic scales is urgently needed to forecast species’ future range shifts and their associated consequences for global biodiversity patterns. Species distribution models (SDMs) are the primary tool used to make these assessments and forecasts. SDMs apply associations between the known occurrences of individual species and co-incident environmental variables, such as temperature and precipitation, to model current species distributions and to forecast future ones^5–7^.

Despite their widespread use and ease of application for more than two decades, most SDMs are based solely on abiotic factors (i.e., the “fundamental” niche, *sensu* Hutchinson’s^8^). Such abiotic SDMs thus ignore important traits and biotic interactions that modulate complex processes influencing species persistence and fitness^9–12^. Although some studies have incorporated biotic traits into SDMs^13, 14^, such “trait-based” SDMs have not been widely adopted. Moreover, existing approaches that integrate traits into SDMs generally require detailed mechanistic information linking trait variation and fitness, limiting the generality of inferences regarding predicted and forecasted species distribution patterns across space, time, and taxa^15, 16^. Recently developed “trait-based” frameworks that integrate functional traits reflecting the intrinsic properties of focal species do not require the complex parameterization necessary for such process-based models and hold tremendous promise to bring greater biological reality—and thus, greater predictive accuracy—into SDMs^14^. However, to the best of our knowledge, these trait-based SDMs to date have not accommodated intraspecific changes in the expression of these traits resulting from co-incident spatial and temporal shifts in abiotic conditions such as those predicted to occur as the climate changes in the near future.

To improve the generality of trait-based SDMs and accommodate climate-driven changes in trait expression, we developed a new type of trait-based SDM that directly incorporates responses of traits to climate change. Specifically, our method simultaneously models known historical data linking traits, climate, and species distributions to forecast where and when climatically suitable areas will exist for focal species (Fig. 1). We tested the utility of our model using phenology—the timing of recurrent life-history events such as leaf-out and senescence, budding, flowering, fruiting, and seed dispersal. Phenology is a key functional trait that responds to climate and ongoing climate change, determines the exposure of reproductive structures to potentially stressful abiotic conditions, influences the ability of a species to capture resources, and affects its interactions with other species^17, 18^. Moreover, phenology is frequently determined by the response of critical life-history stages to varying climatic conditions, such as advancing or delaying the onset of growth and development under warmer versus colder spring temperatures. Phenology has also been demonstrated to be a major component of plant fitness^19^, and can be the primary factor limiting species’ distributions in some ecosystems^16, 20–22^. Phenology also can exhibit substantial intraspecific variability as a function of climatic variation across a species’ range^23–25^. Such intraspecific variability remains unexplored for most species but may confer important resilience to changing climates^9, 22, 26^.

**Fig. 1.**
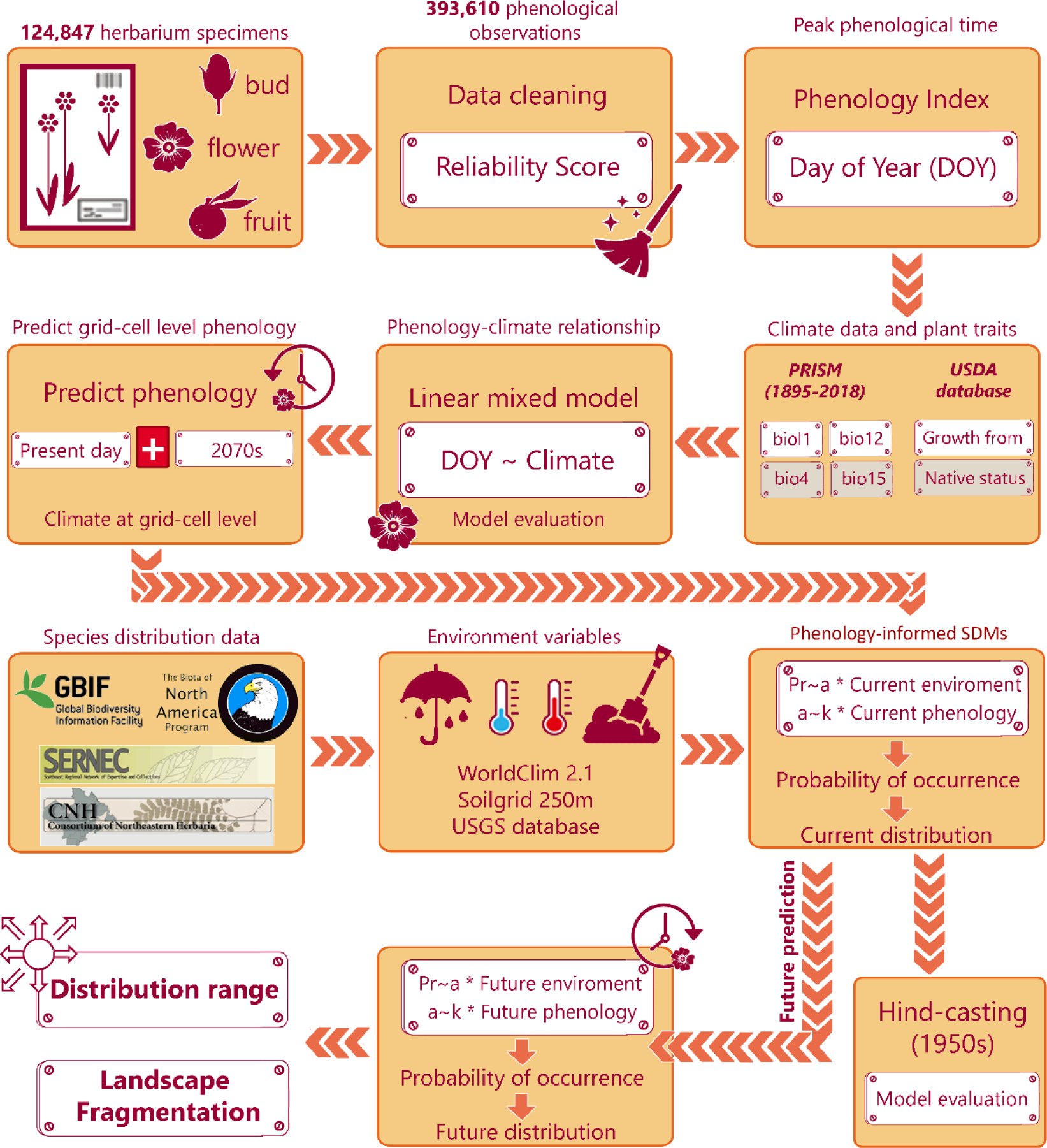
Phenology-informed SDM workflow. Predicted phenology [peak budding, flowering, and fruiting time (Day of Year, DOY)], ascertained from > 120,000 herbarium specimens and ∼400,000 phenological observations under both current and future climates, were estimated for each species per grid cell based on species-specific phenology-climate relationships. We then built phenology-informed SDMs to demonstrate whether phenology mediates responses of individual plants to environmental gradients, and forecasted species distributions and regional diversity under future climate scenarios. To validate model reliability, we hind-casted our model using historical data to project species distributions before 1950 and compared them with actual species distributions recorded before 1950.

To build our new “phenology-informed” SDM, we constructed a large phenological data set of over 120,000 herbarium specimens that includes nearly 400,000 phenological observations (i.e., of buds, flowers, and fruits) from 360 plant species growing in the eastern United States (i.e., from Maine to Florida and westward to West Virginia and Georgia). We then used this phenology-informed SDM to predict changes in species distributions and regional biodiversity as a function of future climate-change scenarios. Our overarching goal was to create a framework for a more dynamic SDM forecast that directly and dynamically integrates phenological responses to climatic changes (**Fig. 1**). Specifically, we i) modeled relationships between climate and phenology to estimate the phenological responses of individual species to varying climatic factors; ii) validated the modeling approach using hindcasting; iii) forecasted the distribution of each species under current and future environmental conditions; and iv) compared the results of our phenology-informed SDM with results from standard abiotic-only SDMs. The results illustrate whether and how phenology shapes species’ distributions along environmental gradients, and how phenology may modulate species’ responses to ongoing and future climate change. Importantly, our novel phenology-informed SDM exemplifies a more expansive trait-based SDM framework that is widely applicable to other fitness-related functional traits that covary with abiotic factors across space, time, and taxa.

## Results

### Plant phenological responses along climatic gradients

Key predictors of day of year (DOY, see Methods) of peak budding, flowering, and fruiting time included annual mean temperature (bio1), temperature seasonality (bio4), precipitation seasonality (bio15), and species’ growth form (Table S1). The strongest climatic predictors for budding, flowering, and fruiting time were bio1 and bio4, but their effects differed significantly among species with different growth forms (Table S1). For example, holding other predictors constant, a 1-standard deviation (SD) increase in temperature (*ca*. 5.3°C) would advance the budding time of herbaceous annuals by 8 ± 3.6 days (mean ± one standard error of the mean [SE]), of herbaceous perennials by 10 ± 3.2 days, and of woody plants by 10.0 ± 3.9 days.

### Parameterizing phenologically-informed SDMs

We first classified all BIOCLIM climate (bio1–bio19) and soil variables into four categories (Table S2; Methods): (1) mean temperature (bio1, bio5, bio6, bio8, bio9, bio10, bio11); (2) mean precipitation (bio12, bio13, bio14, bio16, bio17, bio18, bio19); (3) climate fluctuation (bio2, bio3, bio4, bio7, bio15), and (4) soil (five variables). We then conducted a principal component analysis (PCA) on per-cell values of each environmental group. The first PC of the mean temperature variables accounted for 86% of the variance in mean temperature; the first two PCs of the mean precipitation variables separately accounted for 55% and 39% of the variance in mean precipitation; the two climate seasonality PCs accounted for 72% and 22%, respectively, of the variance in climatic fluctuation; and the first two soil PCs accounted, respectively, for 60% and 32% of the variance in soil composition and structure (see Methods for analytical details of the PCAs and Fig. S1 for detailed results of the PCAs).

All generalized linear mixed models (GLMMs) of plant distributions performed well, with an average explained deviance of more than 90% (conditional *R*^2^) and AUC of 0.932 (Table 1). Plant reproductive phenology had a strong modulating influence on the occurrence of plants along environmental gradients (i.e., significant interactions between phenology and environmental factors; Table 1), and the impacts of phenology on the responses of species to environmental conditions varied among different phenophases and environmental variables (Fig. 2; Table 1). Positive *Z*-statistics suggested that a high value for DOY increased the probability of occurrence of species under high values of the certain environmental variable.

**Fig. 2.**
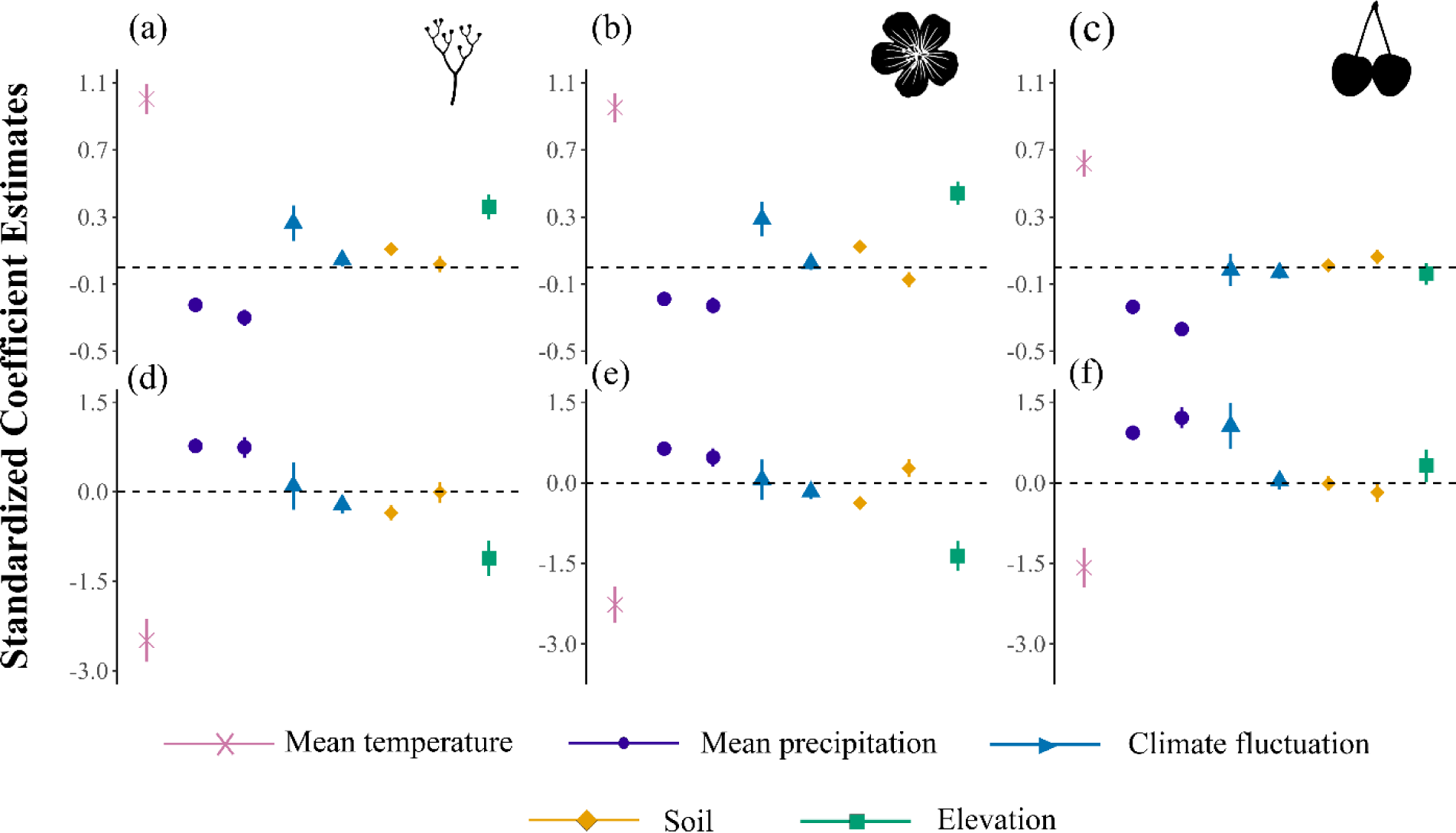
The standardized coefficient estimates indicate the main impacts of different environmental factors on species’ probability of occurrence (a–c) and how phenology mediates average species response to environmental gradients or, alternatively, how environmental effects on the average probability of occurrence are influenced by phenology (i.e., interaction terms between plant phenology and environmental conditions; d–f). We examined the effects on the probability of occurrence of the interaction between phenology and each of five classes of environmental attributes (mean temperature [one PC], precipitation [two PCs], climate fluctuation [two PCs], soil composition and structure [two PCs], and elevation [m asl]). Phenology was quantified as the day of year (DOY) for plant peak budding (a, d), flowering (b, e) and fruiting (c, f). Points and range bars represent among-species average standardized coefficient estimates with 95% confidence intervals (CIs). Standardized coefficient estimates were considered significant if their 95% CIs did not overlap zero.

**Table 1.** Summary of phenology-informed species distribution models (GLMMs). The various principal components (PCs) represent different environmental components and DOY refers to the Day of Year for peak budding, flowering, or fruiting time. The “:” symbol in the phenological traits column represents an interaction between variables.

| Phenological traits | Budding |  | Flowering |  | Fruiting |  |
| --- | --- | --- | --- | --- | --- | --- |
|  | Z | p | Z | p | Z | p |
| Temperature PC1 | -13.525 | < 0.001 | -13.076 | < 0.001 | -8.324 | < 0.001 |
| Precipitation PC1 | 12.431 | < 0.001 | 11.066 | < 0.001 | 14.171 | < 0.001 |
| Precipitation PC2 | 8.397 | < 0.001 | 5.608 | < 0.001 | 12.567 | < 0.001 |
| Climate Fluctuation PC1 | 0.478 | 0.633 | 0.315 | 0.753 | 4.877 | < 0.001 |
| Climate Fluctuation PC2 | -2.829 | 0.005 | -2.250 | 0.024 | 0.486 | 0.627 |
| Soil PC1 | -5.392 | < 0.001 | -6.171 | < 0.001 | -0.187 | 0.851 |
| Soil PC2 | -0.098 | 0.922 | 3.243 | 0.42 | -1.885 | 0.059 |
| Elevation | -7.348 | < 0.001 | -9.643 | 0.001 | 2.092 | 0.036 |
| DOY | 4.162 | < 0.001 | -1.162 | 0.245 | -0.982 | 0.326 |
| Temp PC1: DOY | 21.981 | < 0.001 | 21.298 | < 0.001 | 15.164 | < 0.001 |
| Precip PC1: DOY | -13.111 | < 0.001 | -11.564 | < 0.001 | -15.235 | < 0.001 |
| Precip PC2: DOY | -12.352 | < 0.001 | -9.558 | < 0.001 | -16.507 | < 0.001 |
| Clim Fluc PC1: DOY | 4.848 | < 0.001 | 5.483 | < 0.001 | -0.167 | 0.87 |
| Clim Fluc PC2: DOY | 2.115 | 0.03 | 1.175 | 0.24 | -1.617 | 0.11 |
| Soil PC1: DOY | 6.324 | < 0.001 | 7.582 | < 0.001 | 0.82 | 0.41 |
| Soil PC2: DOY | 0.751 | 0.452 | -3.210 | 0.001 | 2.896 | 0.004 |
| Elevation: DOY | 9.461 | < 0.001 | 12.365 | < 0.001 | -1.03 | 0.30 |
| Conditional R<sup>2</sup> | 0.917 |  | 0.912 |  | 0.913 |  |
| Marginal R<sup>2</sup> | 0.082 |  | 0.079 |  | 0.067 |  |
| AUC | 0.917 |  | 0.946 |  | 0.946 |  |

The probability of occurrence within 40×40-km^2^ grid cells (see Methods) of individual plants with early-season budding, flowering, and fruiting times decreased much more rapidly with increasing temperature than those with later phenology. However, the probability of occurrence of individual plants with earlier budding, flowering, and fruiting times increased more rapidly with increasing precipitation than those with later phenology. Individual plants with late budding and flowering time were more likely to occur in areas with high temperature seasonality but not in regions with high precipitation seasonality. The effects of soil characteristics on plant distributions also varied among individuals with different phenological states. The negative relationship between the probability of occurrence and elevation was much stronger for early budding and flowering individuals than for those with later phenology (Table 1, Fig. 2).

### Hindcast validation of the phenology-informed SDM

Similar to standard abiotic SDMs, our phenology-informed SDMs exhibited good hindcasting performance: an average of 74.5% and 73.8% of the distribution points of 57 annual herbs before 1950 fell within their predicted past range based on the phenology-informed and abiotic SDMs, respectively (Fig. S2).

### Species distributions under future climate change

#### Projected range changes

There were no obvious phylogenetic signals in the proportional change of suitable areas forecasted by both phenology-informed and abiotic SDMs (Table S3), so our models did not include a phylogenetic correction. Using the phenology-informed SDMs, we forecasted that ≈ 35% of the current occupied ranges of species would have low future suitability (range retreat), and ≈ 26% of the species would lose at least half of their existing suitable habitats in the future. In contrast, the abiotic SDMs forecasted that ≈ 40% of the current occupied ranges would have low future suitability, and that ≈ 35% species would lose at least half of their existing habitats. Abiotic SDMs and phenology-informed SDMs showed similar estimates of the species’ current range (paired *t*-test; p = 0.11) but significantly different estimates of the species’ future range. Specifically, the average proportion of current areas forecasted to have high suitability (i.e., range persistence) based on the phenology-informed SDMs was *ca*. 5% higher than that predicted by the abiotic SDMs. In contrast, phenology-informed SDMs forecasted a 20% lower proportion of future occupying regions located outside their current range (i.e., range expansion) than abiotic SDMs (Fig. 3; both differences significant at *p* < 0.05). On average, landscape fragmentation of future species distributions was forecasted to be 30% higher (*p* < 0.05) than that of current species distributions (Fig. S3) by both phenology-informed SDMs and abiotic SDMs.

**Fig. 3.**
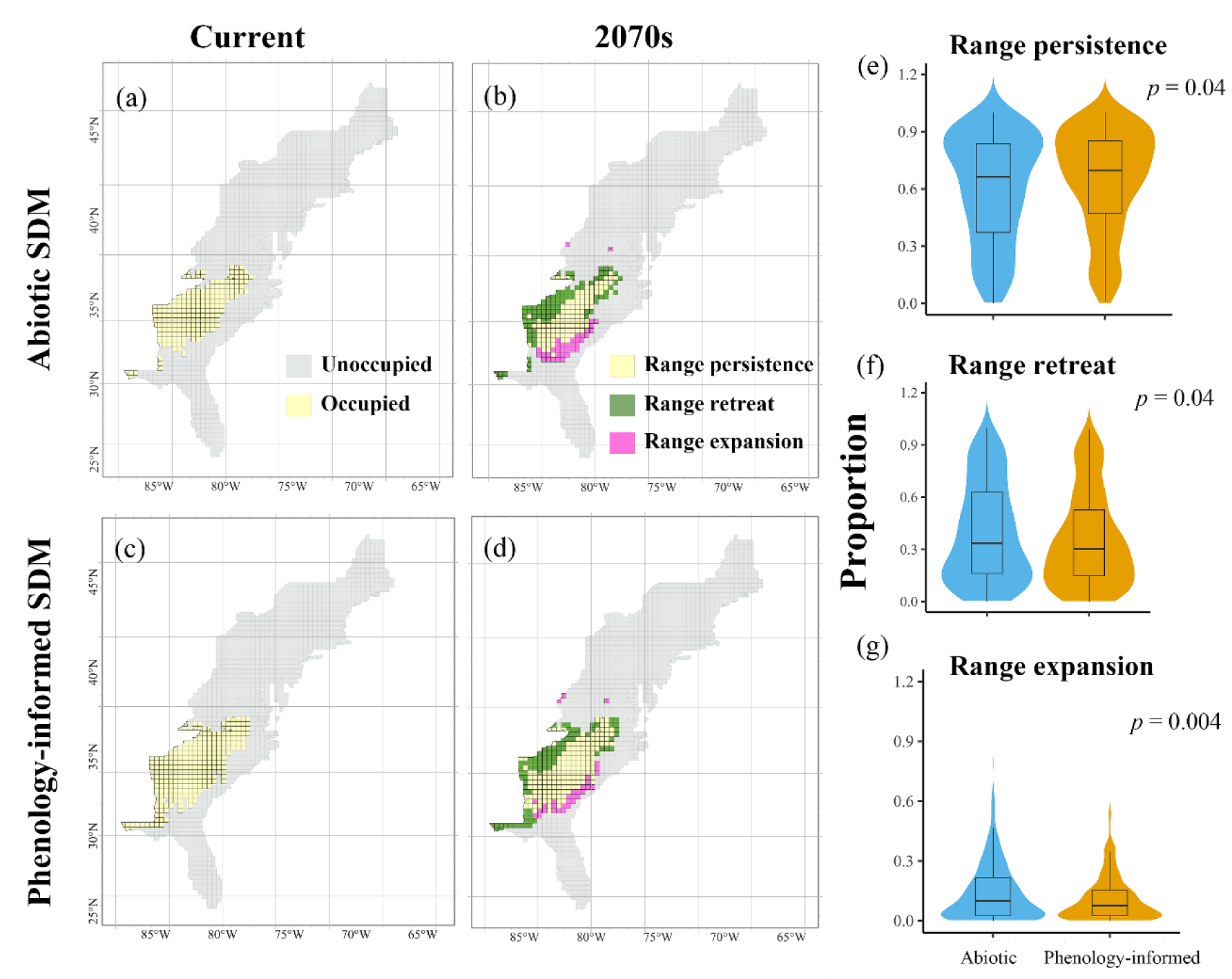
Changes in species ranges forecasted by standard abiotic SDMs and phenology-informed SDMs. Maps illustrate the current (a, c) and future (b, d) distributions of *Trillium catesbaei* as an examplar. Yellow areas represent regions currently occupied by *T. catesbaei* that are projected to remain suitable in the future (i.e., range persistence); green areas represent regions currently occupied by *T. catesbaei* that are projected to have low suitability in the future (i.e., range retreat); pink areas represent regions currently unoccupied by *T. catesbaei* that are projected to have high suitability in the future (i.e., range expansion). Violin plots illustrate the proportion of species’ ranges that persist (d), contract (e), or expand (f) across all 360 species forecasted by abiotic SDMs (blue) versus phenology-informed SDMs (orange). The differences between the two models were evaluated by a paired Student *t-test* (*p* < 0.05).

#### Geographic trends in regional species diversity

At the grid cell-level, we calculated several metrics, including species gains, losses, and turnover, that are commonly used in modeling future biodiversity scenarios. Spatial patterns in absolute species gains and losses were consistent across GCM scenarios (Figs. S4–S6). In general, more gains in species were expected in Florida and the Atlantic coastal plain than in New England and parts of the Appalachian Mountains (Fig. 4d, e; Fig. S7). The forecasted percentage of species losses exceeded 50% in the southern Coastal Plains, some portions of the Atlantic coastal plain, and the southern extent of the Appalachian Mountains. The combination of high species gains and high species losses in Florida and the Atlantic Coastal Plain were forecasted to lead to high turnover rate in these regions (Fig. 4f; Fig. S7). We identified significant linear relationships among grid cells between the percentage of absolute species gains (and losses) and the magnitude of change in mean annual temperature. Relative to the abiotic SDMs, the phenology-informed SDMs predicted relatively lower species gains when temperature was lower than 5 °C but higher species gains at higher temperature (Fig. 4a). However, phenology-informed SDMs generally predicted stronger negative relationships between species losses (and turnover) and mean annual temperature than abiotic SDMs (Fig. 4b, c)

**Fig. 4.**
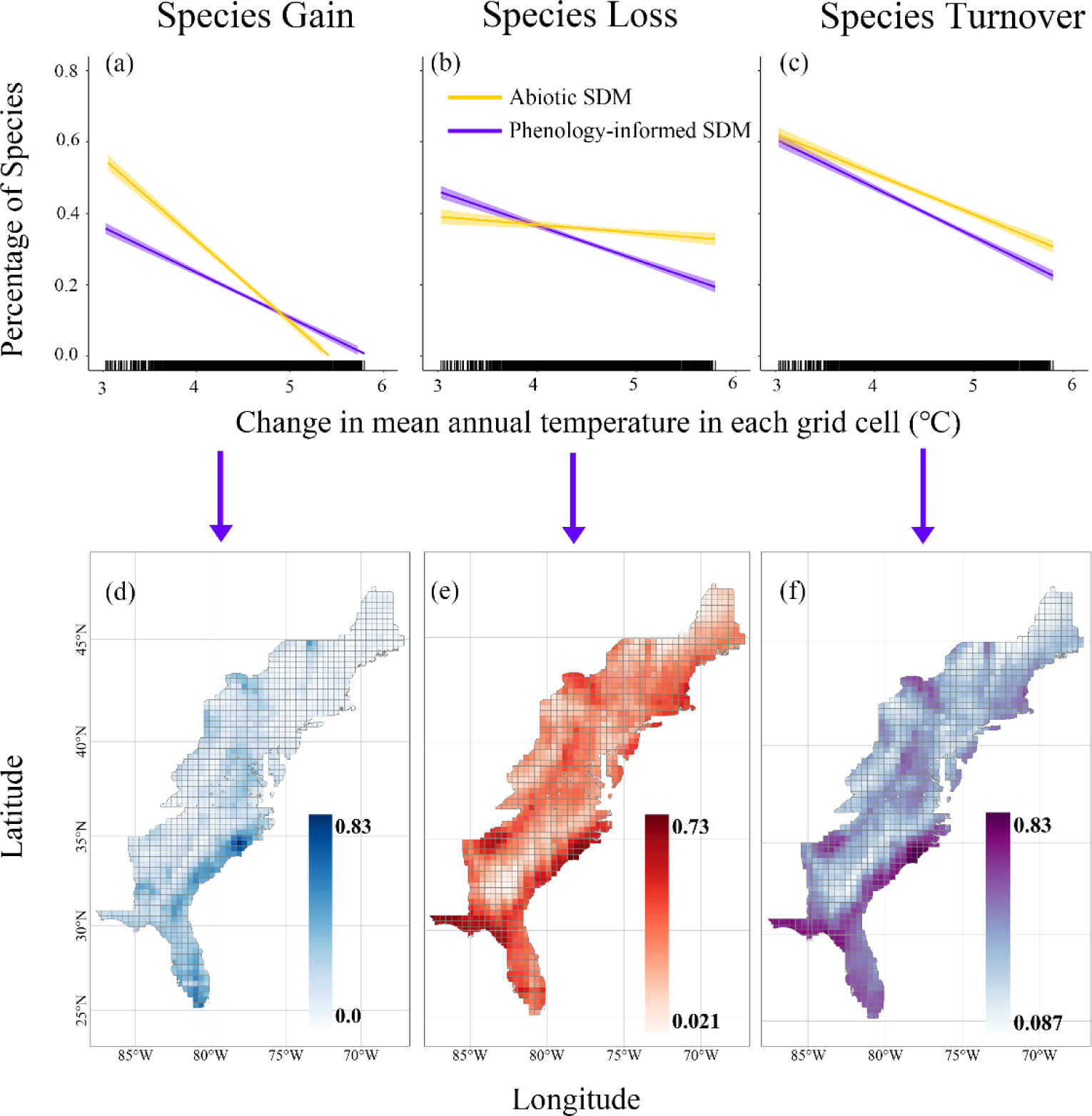
Changes in species composition within each grid cell under future climate change. (a–c) Relationships between the proportion of species gains, losses, and turnover and changes in mean annual temperature within each individual grid cell. These relationships were separately explored by abiotic SDMs and phenology-informed SDMs. Here, phenology was quantified as the day of year for peak flowering, and mean annual temperature in each grid cell was calculated as the average across all six General Circulation Models under the SSP5-85 climate scenario in the 2070s. (d–f) Geographical patterns in the proportion of species gains, losses, and turnover forecasted by phenology-informed SDMs.

## Discussion

SDMs are widely used to assess the potential consequences of climate change for species’ distributions^27^. However, these models have been criticized for omitting critical biological components, including intraspecific phenotypic variation in functional traits, which may have important consequences for species fitness^28^ or for the ability of a population to persist at a given location. Our newly developed phenology-informed SDM incorporates key functional trait data to explore how changes in phenology affect forecasted species geographic distributions in the face of climate change. In general, both abiotic and phenology-informed SDMs forecast an increase in fragmentation of, and long-term decline in, species’ ranges in the eastern United States. However, our model inferences differed in one key and important way: our phenology-informed SDM inferred from decades of herbarium specimens suggested a more optimistic forecast for species distributions than did the more simplistic and commonly used abiotic SDMs.

Our approach differs from existing process-based models that integrate the impacts of climate-mediated phenology on demography and fitness to estimate species distributions^16, 20^. Specifically, these methods require detailed knowledge of the mechanistic basis of phenology and local dynamics within a system to implement effectively. In contrast, our method does not require such knowledge and thus can enable scaling range-shift assessments to include thousands of species by using existing taxonomically and geographically extensive resources (e.g., herbarium specimens). Our approach also differs from the method used by Hereford et al.^21^, who constructed a seasonal SDM for the very short-lived, ephemeral, *Mollugo verticiillata* using herbarium records. Hereford et al.^21^ used the localities and dates of collections, along with monthly climate data, as proxies of phenology to map the future distribution of this species for each month of the year. In contrast, our method simultaneously uses direct phenological information alongside trait-climate relationships, which also explicitly account for the co-variation between reproductive phenology and climate across the range of each species. Further, our approach is generalizable to other fitness-related traits amenable to a trait-based SDM.

### Phenology mediates the effects of climate on species’ distributions

We identified substantial variation in the predicted peak budding, flowering, and fruiting time among individual species based on phenology (DOY)-climate relationships. Our estimates of phenological responses to climate parallel those we reported previously for the eastern United States that demonstrated broad and variable phenological responses within and among species^24^. By accounting for phenology-climate relationships in our SDMs, we demonstrate that phenology can modulate the effects of climate on plant occurrence across broad environmental gradients (Fig. 2; Table 1). This suggests that phenological variation accounts for substantial variability in the ability of individual plants to withstand climate change.

Specifically, the probability of occurrence of individual plants that reproduced relatively early in the growing-season decreased more with temperature and increased more with precipitation than plants that flowered relatively late in the growing season (Fig. 2; Table 1). Different life-history stages (e.g., growth, reproduction) are biologically linked and the timing of these phases can have direct bearing on reproductive allocation per offspring. For example, in temperate climates, large-seeded species often flower earlier than small-seeded species, presumably because the former require more time to develop and ripen their seeds^29, 30^. In contrast, late-flowering species often allocate more resources to maternal growth, but the time available for seed development may be decreased due to the ending of the growing season^31^. Large seeds subsequently produce larger seedlings, some of which are more drought resistant during the late summer and early fall^32^. On the other hand, later-flowering plants often are exposed to risks associated with early frosts during seed maturation^33^. From this perspective, late-flowering plants may be more favored under warming scenarios in which the growing season is extended^33, 34^. In support of these hypotheses, we identified higher probabilities of occurrence of late-flowering plants than those of early-flowering plants at higher elevations (Table 1). Higher elevations tend to be colder and have shorter growing seasons than lower elevations. Plants growing at higher elevations usually have lower seed production, which could be offset by vegetative growth during longer seasonal vegetative periods (i.e., late flowering) to ensure the persistence of populations^35^.

### Phenology-informed SDMs forecast less severe habitat loss of species in response to climate change

Climate change is contributing to widespread range contractions and local extirpations of species^36, 37^. Urban^27^ projected that approximately one in six species will face extinction if climate change proceeds as expected. Our results predict increased species geographic range fragmentation and species geographic range losses for both phenology-informed and abiotic SDMs. Both models forecasted that > 50% of the species we sampled will experience both range loss and increased habitat fragmentation under future climate change scenarios. However, our results differed between these models in important ways.

First, although the degree of inferred fragmentation was forecasted to increase in the future (and to similar extents) by both phenology-informed and abiotic SDMs (Fig. S3), changes in the sampled species’ geographic ranges differed significantly between the two types of SDMs. The abiotic SDMs predicted that, on average, across species, 60% of the currently occupied ranges of species would have high suitability in the future, whereas the phenology-informed SDMs predicted that an average of 65% of the currently occupied ranges would persist (Fig. 3). These findings provide evidence that the geographic range of a species is modulated not only by climate but also by variation in functionally relevant biological traits that allow populations to survive within their local environments^38^. As a vital fitness-related trait, plant reproductive phenology exhibits high intraspecific variation in phenological sensitivity across species’ ranges^24^. Thus, the fundamental assumption of abiotic SDMs—homogeneous responses to climate variations across their range—is unrealistic. SDMs that do not include information on traits and their dynamic interactions with the environment and with other traits may result in incorrectly estimating a species’ climatic niche, which may, in turn, lead to incorrect forecasts of species loss.

As the climate continues to change, populations are expected to persist *in situ* via local adaptation and phenotypic plasticity, track climate through migration, or become locally extinct^39^. The first two biological mechanisms are important for the persistence of species by expanding the climatic tolerance of species beyond their present realized niches.

Species that can maintain their climatic niche by acclimating their phenology to changing climates may not need to migrate to survive and reproduce^40^. Therefore, climate-induced phenological responses may alter expectations of species’ distributions under future climatic conditions, a prediction consistent with our phenology-informed SDMs, which generally predict a smaller reduction in species’ ranges with climate change compared to forecasts generated by abiotic SDMs.

### Phenology-informed SDMs predict less species turnover in response to climate change

We found significant geographic differences in species gains, losses, and turnover. Florida and the Atlantic Coastal Plain were forecasted to have a higher proportion of species losses. We hypothesize that this pattern may be driven by two factors: a narrower range of phenological response to temperature among southern populations and a thermal tolerance maximum reached by these populations under projected climate change. Although the absolute amount of warming is lower at low latitudes, many species with narrow ranges are endemic to the southeastern U.S. and the southern Appalachian Mountains. These species frequently exhibit more narrow climatic tolerances, which are likely to promote high rates of species loss under future climate change^41^. These species may also be nearer to their thermal maxima, which may be exceeded in the southeastern United States in forecast non-analog climatic conditions^42^. We additionally found a significant positive relationship between the mean degree of warming predicted across the range of species and the proportion of range that persists (Fig. S8), which supported our results of high species loss in southern regions with a relatively low magnitude of warming compared to northern regions.

We identified strong negative relationships between the percentage of species gains, losses, and turnover and the expected amount of warming for individual grid cells. Qualitative patterns were similar for both phenology-informed and abiotic SDMs. However, phenology-informed SDMs generally forecasted a lower proportion of species losses and turnover in response to warming. We hypothesize that both phenotypic plasticity and adaptive components of plant phenology contribute to persistence of populations as the climate changes and tend to increase opportunities for species migration to the extent that they contribute to species niche breadth^43, 44^. Our phenology-informed SDMs forecast more persistent regional species composition under climate warming relative to the abiotic SDMs, and we suggest that previous studies using only climatic variables in SDMs may overestimate the impacts of climate change on species turnover.

In summary, both abiotic and phenology-informed SDMs project species fragmentation and range loss across hundreds of plant species in the eastern United States under future climatic change scenarios. However, phenology-informed SDMs forecast significantly less drastic species’ range loss and turnover within communities. We conclude that future research and conservation efforts should look beyond abiotic SDMs and embrace and integrate biological phenomena that contribute to species-specific acclimation and adaptive responses to climate change. Such work also may reveal climatic tolerances beyond those predicted by standard abiotic SDMs. Finally, whereas our study uses phenology as an exemplar trait with which to build taxonomically and geographically broad SDMs, our methodological framework could be further developed to explore how other climate-sensitive traits respond. Obvious traits to explore include photosynthetic modality (e.g., C4 versus C3 plants), symbiotic associations (e.g., seed and fruit dispersers, pollinators, and hosts), and dispersal ability.

## Methods

### Species selection and phenological data collection

We used digitized specimens from two of the most comprehensive digitized regional floras in the world: the Consortium of Northeastern Herbaria (CNH, https://neherbaria.org/portal/) and the Southeast Regional Network of Expertise and Collections (SERNEC, https://sernecportal.org/portal/). These two online portals include more than six million digitized herbarium specimens from across the eastern United States. Following previously established protocols^24, 45^, we selected species and specimens for analysis when i) there were at least 50 unique collections of the species across space and time in the eastern United States, ii) the specimens included both an exact collection date and at least county-level location information, and iii) the specimens had easily identifiable and quantifiable buds, flowers, and fruits. Applying these criteria yielded 124,847 total herbarium specimens representing 360 species that we used in our models and analyses. These species vary in life history, growth form and native status (i.e., native or not to the eastern United States) (Table S4). Information on species’ growth form and native status were derived from the United States Department of Agriculture PLANTS Database (https://plants.isda.gov/). Our final database included 57 herbaceous annuals, 224 herbaceous perennials, 12 vine perennials, 67 woody species, of which 314 are native and 46 are introduced.

### Phenological data extraction

Crowdsourcers hired through Amazon’s Mechanical Turk service (MTurk; https://www.mturk.com/) counted the number of each type of structure (i.e., buds, flowers, and fruits) present on the specimen using the *CrowdCurio* platform following the protocols outlined by Willis et al.^42^. Each specimen was scored independently by at least three crowdsourcers. We estimated the reliability of each crowdsourcer and their data following Williams et al.^42^ and Park et al.^24, 45^. Specifically, each 10-image set scored by one person included nine unique images and one duplicate image randomly selected from the other nine. The nine images plus the duplicate image were presented to each crowdsourcer in a random order. We then calculated a reliability score for each crowdsourcer based on the data for each 10-image set. The absolute differences between the two duplicate specimens in counts of buds, flowers, or fruits, were separately divided by the total counts for each phenological state, and then subtracted from 1 (Equation1):

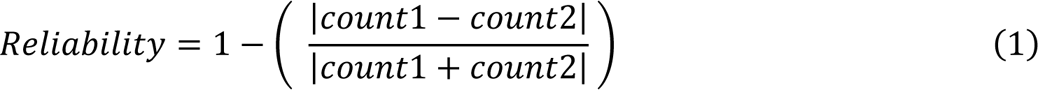

The reliability score ranged from 0 (unreliable) to 1 (reliable). Specimen observations scored by crowdsourcers with a reliability score of zero were excluded from the analysis^24^; If an individual received a reliability score equal to zero for one phenological phase, the reliability scores for that same individual’s counts of the other phenological states were also set to zero (and similarly excluded from the analyses). Our final dataset included a total of 393,610 phenological observations spanning 124 years (1895–2018), of which 157,435 observations have already been published^24, 45^, and 236,175 observations were new to this study.

We calculated the median number of buds, flowers, and fruits from the multiple phenological observations for each specimen. We then estimated the day of year (DOY) of “peak” budding, flowering, and fruiting if their buds, open flowers, or fruits separately represented ≥ 50% of the total number of reproductive structures (as in Park et al.^45^). Two species each were excluded from the “peak budding” category (*Primula mistassinica* and *Sarracenia rubra*) and the “peak fruiting” category (*Calopogon barbatus* and *Narcissus poeticus*) because there were no specimens of these species at these developmental stages. Of the original 124,847 herbarium specimens, 29,094 specimens were scored as being at peak budding (representing 358 species), 55,767 specimens at peak flowering (representing 360 species), and 37,071 specimens at peak fruiting (representing 358 species).

### Species distribution data

County-level species’ distribution data were obtained from the Biota of North America Program’s (BONAP; http://www.bonap.org/) North American Plant Atlas (NAPA; Kartesz, 2015), which included 19,039 taxa from 227 families (accessed November 2022). These data are available as binary occurrences (i.e., presence/absence) for 3,067 counties in the USA, excluding Alaska and Hawai’i. We supplemented the species’ distribution records from BONAP using additional data from the GBIF database (https://www.gbif.org) and all available specimen records in the CNH and SERNEC database. Most (≈ 70%) of these additional data points (i.e., GBIF+NH+SERNEC) represented specimens collected between 1960 and 2000.

Since the median area of all counties across the eastern United States is 1221 km^2^, we divided the contiguous eastern United States into 40 × 40-km (1600-km^2^) grid cells using the Mercator projection (EPSG 3857). After removing incomplete grid cells (those < 800 km^2^) within national borders and coasts, a total of 1,158 grid cells remained. We then overlaid grid cells with county-layer species-occurrence data. A species was deemed “present” in a grid cell if more than half of the cell’s area was covered by the county-level distribution of that species. If a grid cell included parts of several counties in which a species occurred, the total coverage of this grid cell was defined as the sum of all intersected parts of those counties.

### Environmental data

#### Climatic data of specimen localities

We extracted average monthly temperature and precipitation data (1895–2018) at a 4-km resolution from PRISM (product AN81m, http://prism.oregonstate.edu/) to characterize the climate of the locality where and when each specimen was collected. Following Park et al.^45^, we used the climatic data for the county centroid if precise coordinate data were not available for historical specimen records. Parks^46^ found that within-county climatic variation had little effect on estimates of phenological response in the eastern United States. Based on the extracted monthly temperature and precipitation data, we estimated annual mean temperature (bio1), temperature seasonality (bio4), annual precipitation (bio12), and precipitation seasonality (bio15) for each locality-year combination and assigned these values to corresponding specimens. These climatic variables have been shown to be strongly associated with plant budding, flowering, and fruiting times^24, 45, 47^.

#### Environmental data used for modeling species’ distributions

“Recent” (1970–2000) and future (2061–2080; henceforth referred to as “2070s’’) climatic data at 2.5-arc-minute (*ca*. 5-km) resolutions were obtained from WorldClim (https://www.worldclim.org/, *ver.* 2.1; all 19 climatic variables: bio1–bio19; see Table S2). Elevation data with a spatial resolution of 30-arc-seconds (*ca*. 1-km) were obtained from the US Geological Survey. We also included five soil variables—sand content, clay content, silt content, bulk density, and coarse fragments— because previous studies have demonstrated that soil structure may improve the fit of SDMs^48, 49^. We calculated the mean values of each of these soil variables at two soil depths (0–5 cm and 5–15 cm) using soil data from the SoilGrids250m database (https://www.soilgrids.org/) and assumed that these values would be constant through time. We did not include soil chemical variables (e.g., soil pH) that can have high temporal variability. The values for climate, soil, and elevation data assigned to each 40 × 40-km grid cell were the means of all data points within it.

Future climatic projections were derived from the General Circulation Models (GCMs) used by the Coupled Model Intercomparison Project Phase 6^50^ (CMIP6) for four Shared Socio-economic Pathways (SSPs). We used the most extreme, SSP5-85 projections, which have similar 2100 radiative forcing levels as its predecessor (i.e., Representative Concentration Pathway; RCP 8.5). Previous studies have demonstrated that current emission trajectories have more closely tracked RCP 8.5 than other RCP scenarios^51^. To account for potential uncertainties in projections induced by different GCMs, we forecasted future species distributions using six different GCMs: ACCESS-CM2, CMCC-ESM2, GISS-E2-1-G, HadGEM3, INM-CM4-8 and MIROC6.

### Phylogenomic data

We tested for the influence of phylogeny as a significant variable in our linear models (LMM) assessing phenological associations with climate. As part of an ongoing investigation exploring the relationship between phylogeny and phenology (unpublished data), we sampled total genomic DNA from well-annotated herbarium specimens for most species included here. We used the PhyloHerb informatic pipeline to assemble, annotate, and analyze these genomic data^52^. Our maximum likelihood phylogeny was inferred from ∼80 plastid genes and a smaller number of nuclear loci (nr ITS, and ETS) using well developed, widely-employed genome skimming methods^53, 54^. Plastid phylogenomics, in particular, has been central to plant macroevolutionary research for the past 30 years and plastomes are being sequenced and aggregated by the hundreds to resolve both broad relationships among seed plants and genetic variation within populations^55, 56^. The phylogeny inferred from these data were included as a covariate in our initial LMMs as described below to assess for phylogenetic conservatism in phenological response to climate.

## Statistical modeling

### Relationships between climate and phenology

We first applied LMMs to all herbarium specimens pooled across the 360 species to identify species-specific relationships between plant phenology and climatic variables. Separate but identically structured models were built for peak budding, flowering, and fruiting. For each full model, we used the DOY for the phenological state recorded for each specimen as the response variable. Predictor variables in the full model included: the year of specimen collection, four climatic variables (bio1, bio4, bio12, and bio15; all centered and scaled), and two species-level plant traits (growth form and native status) and their interactions (bio. * Growth form; bio. * Native status) as fixed terms. The full model also included random intercepts among the species and sampling locations, and random slopes among species for bio1, bio4, bio12, and bio15. The eight interaction terms between native status or growth form and climate indicated whether changes in plant phenology along climatic gradients differed among species with different growth forms and native status. We did not find obvious nonlinear relationships between DOY and any climatic variables in the preliminary analysis for most species, so we did not include quadratic terms in our models.

All models were fitted using the “lmer” function in the “lmerTest” package^57^ (*version* 3.1-2) of the R software system (*version* 4.2.1). We also checked residuals of all models to ensure that all assumptions were met and used Moran’s *I* to confirm that there was no evidence of spatial autocorrelation. No models showed significant spatial autocorrelation (*p* > 0.05). To predict peak budding, flowering, and fruiting time of each current and future species-grid cell combination, we used the “predict” function in the base “stats” package (*version* 4.0.0) with fits of the models previously described and the estimated values of recent and future bio1, bio4, bio12, and bio15 of each grid cell.

Ignoring phylogenetic relationships among species when fitting LMMs may lead to Type I statistical errors^58^. We first tested for phylogenetic signal in the DOY of peak budding, flowering, and fruiting (i.e., conditional modes of intercept) and in their shifts in response to environmental changes (i.e., conditional modes of slope) using both Blomberg’s *K*^59^ and Pagel’s lambda^60^. Although some variables showed significant phylogenetic signals, but only marginally and only for one of the two metrics (Table S5). We further checked our models with phylogenetic LMMs that accounted for phylogenetic relationships among species using the “phyr” package^61^ (*version* 1.0.2). Preliminary results were consistent with the results based on LMMs (Table S6–S8), so the results presented in the main text were reported based on LMMs without phylogenetic correction.

### Constructing phenology-informed SDMs

We quantified how plant peak budding, flowering, and fruiting time (DOY) separately influenced the effects of environmental change on the species’ probability of occurrence. We first divided the 19 WorldClim environmental variables and the five soil variables into four groups (Table S2): (1) mean temperature (bio1, bio5, bio6, bio8, bio9, bio10, and bio11); (2) mean precipitation (bio12, bio13, bio14, bio16, bio17, bio18, and bio19); (3) climatic fluctuation (bio2, bio3, bio4, bio7, and bio15); and (4) soil variables. Principal component analysis (PCA) was then used to reduce the dimensionality of each group using the “prcomp” function from the “stats” package (*version* 4.0.0). In the subsequent analyses, we used either one or two principal component scores (PCs) as predictor variables in our models. For temperature variables, we used only the first PC, which explained 86% of the variation among the seven temperature variables and was positively correlated with all temperature variables (r > 0.8). For the remaining three groups, we used the first two PCs of each group (Fig. S1). Mean precipitation variables loaded positively on the first two PCs for precipitation. Temperature seasonality was positively correlated and precipitation seasonality was negatively correlated on the first two PCs for climatic fluctuation variables. Coarse fragments and silt content were positively correlated with the first two PCs of the soil variables, whereas soil bulk density loaded negatively on these two PCs. We multiplied all of these PC scores by -1 to reverse their signs and improve their interpretability. High values of PCs corresponded to warmer temperatures, more precipitation, larger climatic variance, and greater clay and silt content and coarse fragments (Fig. S1). Elevation was additionally included as a single variable. Our final phenology-informed SDM used elevation, the aforementioned seven PCs and the predicted grid-cell level phenology, which all were weakly correlated (|r| < 0.7; Fig. S9).

We used generalized linear mixed models (GLMMs) as the framework for our phenology-informed SDMs^14^. We fitted GLMMs using restricted maximum likelihood (REML) with the R package “lmer4”^62^(*version* 1.1-27). We fitted a random intercept, random slope binomial model with a logit link function to species × grid cell presence/absence data (Equations 2 – 5).

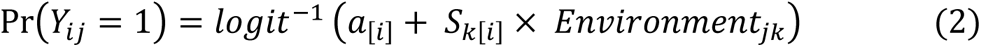

In this base model, the logit probability that species *i* occurs at the *j*th of 1, 2, 3,…,1158 grid cells is equal to an intercept term plus the product of a matrix of eight environmental variables (*Environmental*_*jk*_)and a vector of eight coefficients (*S*_*k*[*i*]_). Here, *k* represents environmental variables. The parameters *a*_[*i*]_ and *S*_*k*[*i*]_ differed among species:

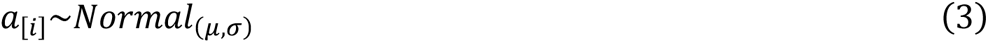

The submodel for *a*_[*i*]_ includes the parameter *μ*, which represents the average probability of occurrence (on a logit scale) among species within a hypothetical grid cell, and the parameter *σ*, which is the degree to which a given species departs from average probability of occurrence. The parameter *S*_*k*[*i*]_ indicates the response of a given species to the relevant environmental variables.

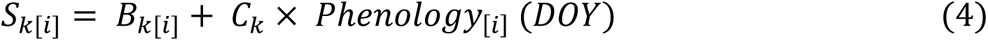

In the submodel for *S*_*k*[*i*]_, its estimate is calculated as the intercept *B*_*k*[*i*]_ plus the coefficient matrix *C*_*k*_ and matrix of trait values *Phenology*_[*i*]_. The intercept *B*_*k*[*i*]_ is modeled as:

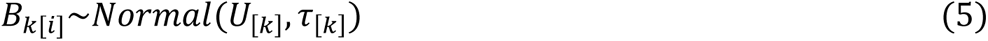

where *U*_[*k*]_ indicates the average response of species to environmental variables and *τ*_[*k*]_ reflects the degree to which each species departs from the among-species average response. *C*_*k*_ describes how plant phenology regulates the response of an individual species to the environmental variables (i.e., DOY * PC_[k]_ interaction coefficients). For example, positive values of *C*_*k*_ suggest that a high value for DOY (i.e., later phenology) increases the probability of occurrence of a species under high values of the focal environmental variable. We also added species as a random component for the slope of occurrence vs. environment, which allowed us to explore species-specific differences in responses to environmental variables^13, 14^. We also constructed abiotic SDMs without phenological information (i.e., only equation 2 and 3) to compare the results of phenology-informed and abiotic SDMs. In abiotic SDMs, we also included species as a random component.

To assess model performance, we calculated the conditional *R*^2^ reflecting the total variance explained^62^. We also calculated the specificity, false negative and false positive rates manually and calculated the area under the curve (AUC) with the “pROC” package^63^ (*version* 1.16.1). AUC represents the predictive accuracy of the model, and ranges from 0.5 to 1 (1 is the highest accuracy). We then converted the probability of occurrence of each species × grid-cell combination into binary maps with the threshold that maximized the specificity and the sensitivity, using the “ROCR” package^64^ (*version* 1.0-11).

### Model validation through hindcasting

To validate our model, we created phenology-informed SDMs using distribution and phenology records of annual herbs (57 species) between 1970-2022, and then hindcasted the models using historic climatic data that originated from PRISM to project species distributions before 1950. We chose annual herbs as a validation group because their short generation times make it likely that rapid evolution has occurred for both phenological plasticity and mean phenological timing in response to environmental fluctuations^65, 66^. In contrast, because of their long lifespans, phenological changes associated with climate change observed for woody plants and perennial herbs are more likely to reflect only phenotypically plastic responses^67^. Our phenological model could not differentiate between climate-mediated phenological variation attributable to phenotypic plasticity or local adaptation. The characteristics of annual herbs are closer to our model assumption. We selected 1950 and 1970 as critical time points because species experienced rapid climate change during the interval from 1950–1970^68, 69^. We compared the projected species distributions with actual species distribution points recorded before 1950.

### Data analyses

We calculated three metrics of changes in suitable areas of species occurrence under future climate change (combining all the SSP5-85 GCMs): i.) current occupied range forecasted to have high suitability (i.e., range persistence; probability of occurrence > threshold); ii.) current occupied range forecasted to have low suitability (i.e., range retreat; probability of occurrence ≤ threshold); and iii.) current unoccupied range forecasted to have high suitability (i.e., range expansion). Present and future suitable areas were determined, respectively, from the number of grid cells where a species occurred now or was forecasted to occur in the 2070s. To examine whether phylogenetic relationships explain the degree of predicted range preservation, expansion and or contraction among species, we calculated the proportion of each species’ range persistence, range retreat, and range expansion and then tested for phylogenetic signal in these three traits using Blomberg’s *K* ^59^ and Pagel’s lambda^60^. Previous studies have suggested that climate change also could lead to fragmentation of species habitats^70^. Based on species distribution maps under current and future conditions, we further calculated the habitat fragmentation index for each species using the “lsm_c_pd*”* function in R package “landscapemetrics”^71^ (*version* 1.4.3).

To assess the percentage of local extinctions for a given grid cell, we then examined the geographical patterns in the proportion of species loss based on the species’ current and future distributions, which was calculated as the number of species lost (*L*) divided by the current species richness (*S*) of each grid cell (*L/S*). The same procedure was also applied to evaluate the proportion of species gained (*G*) in each grid cell under the assumption that species could colonize any new suitable climate space within the eastern United States and that their future distributions will not be limited by dispersal. The proportion of species turnover is then given by *T = (L + G) / (G + S)*. Relationships between the proportion of species gains, losses, and turnover as a function of different degrees of climate change (e.g., changes in mean annual temperature) were further explored to examine the stability of species composition under climate fluctuations. All of these analyses were separately conducted for the abiotic and phenology-informed SDMs. We compared the changes in species’ suitable areas and geographical patterns in species gains (losses and turnover) between abiotic and phenology-informed SDMs.

## Acknowledgements

We acknowledge fundings from the National Science Foundation funding grants: DEB2105903, DEB1754584, EF1208835, DEB2101884, DEB1802209, DEB2105932 and MRA2105903.

## Author Contributions

CCD and AME proposed the initial idea for the study with subsequent development by SP; SP and AME designed the study; SP, CCD, AME collected the data; SP analyzed the data; SP drafted the first version of the manuscript, and all authors contributed significantly to subsequent revisions.

## Ethics declarations

The authors declare no competing interests.

## Data availability

All data used for analyses are publicly available on Github (https://github.com/Shijia818/Phenology-informed-SDM) for the moment and will be archived at the Dryad Digital Repository once accepted.

---

**Fig. S1.**
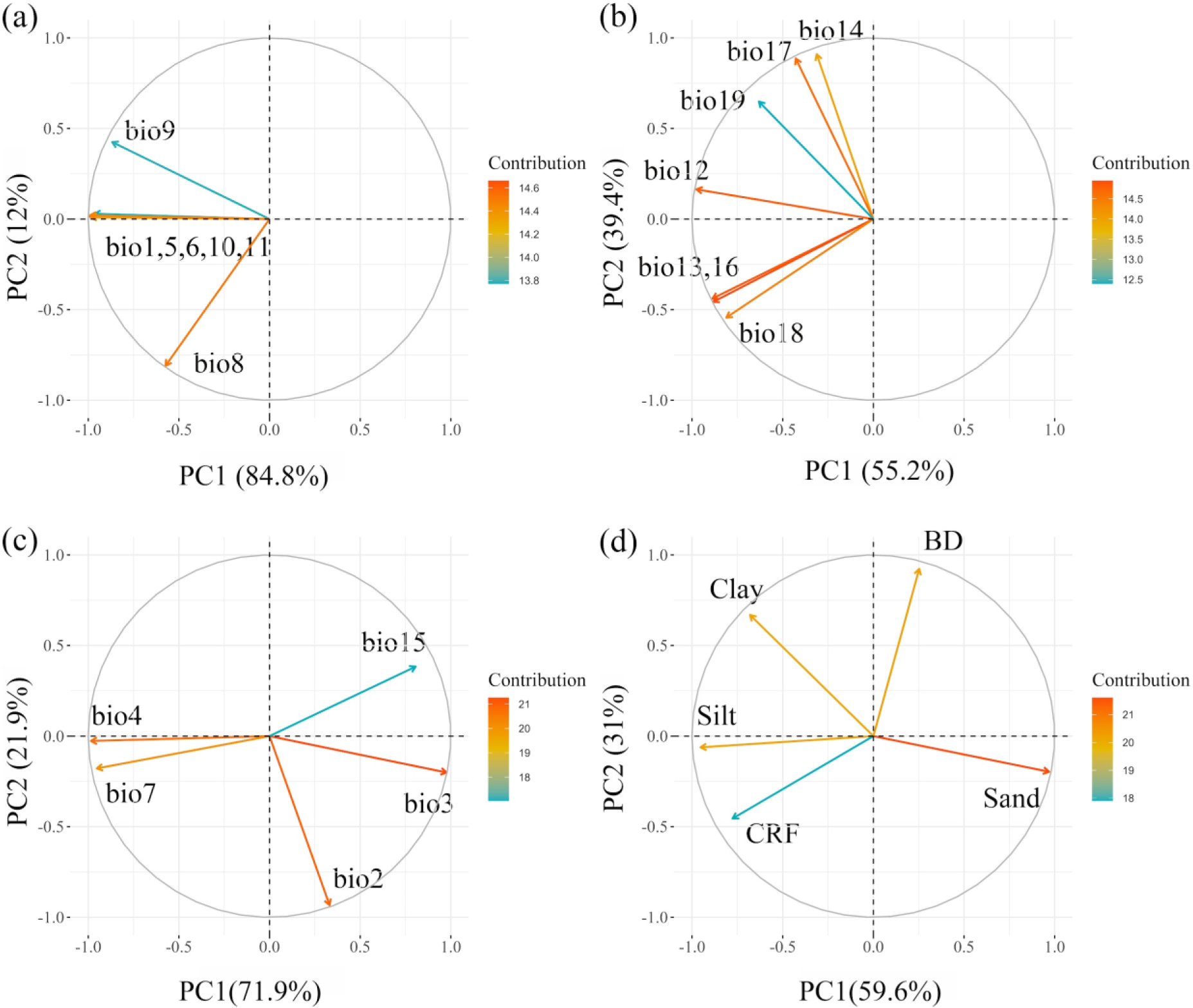
Principal component analysis (PCA) ordination diagram reflecting the contribution of the environmental variables on the first two axes of the PCA. All environmental variables were classified into four groups: mean temperature (a), mean precipitation (b), climatic fluctuation (c) and soil variables (d). Bioclimatic variables (bio1-bio19) were derived from *WorldClim* database (*verison*. 2.1). Abbreviations for soil variables: BD, bulk density (cg/cm^3^); Sand, sand content (g/kg); Clay, clay content (g/kg); Silt, silt content (g/kg); CRF, coarse fragments (cm^3^/dm^3^). PC1 in the main text refers to PC1 of panel (a); PC2 and PC3 in the main text refer to PC1 and PC2 of panel (b); PC4 and PC5 in the main text refer to PC1 and PC2 of panel (c) and PC6 and PC7 in the main text refer to PC1 and PC2 of panel (d).

**Fig. S2.**
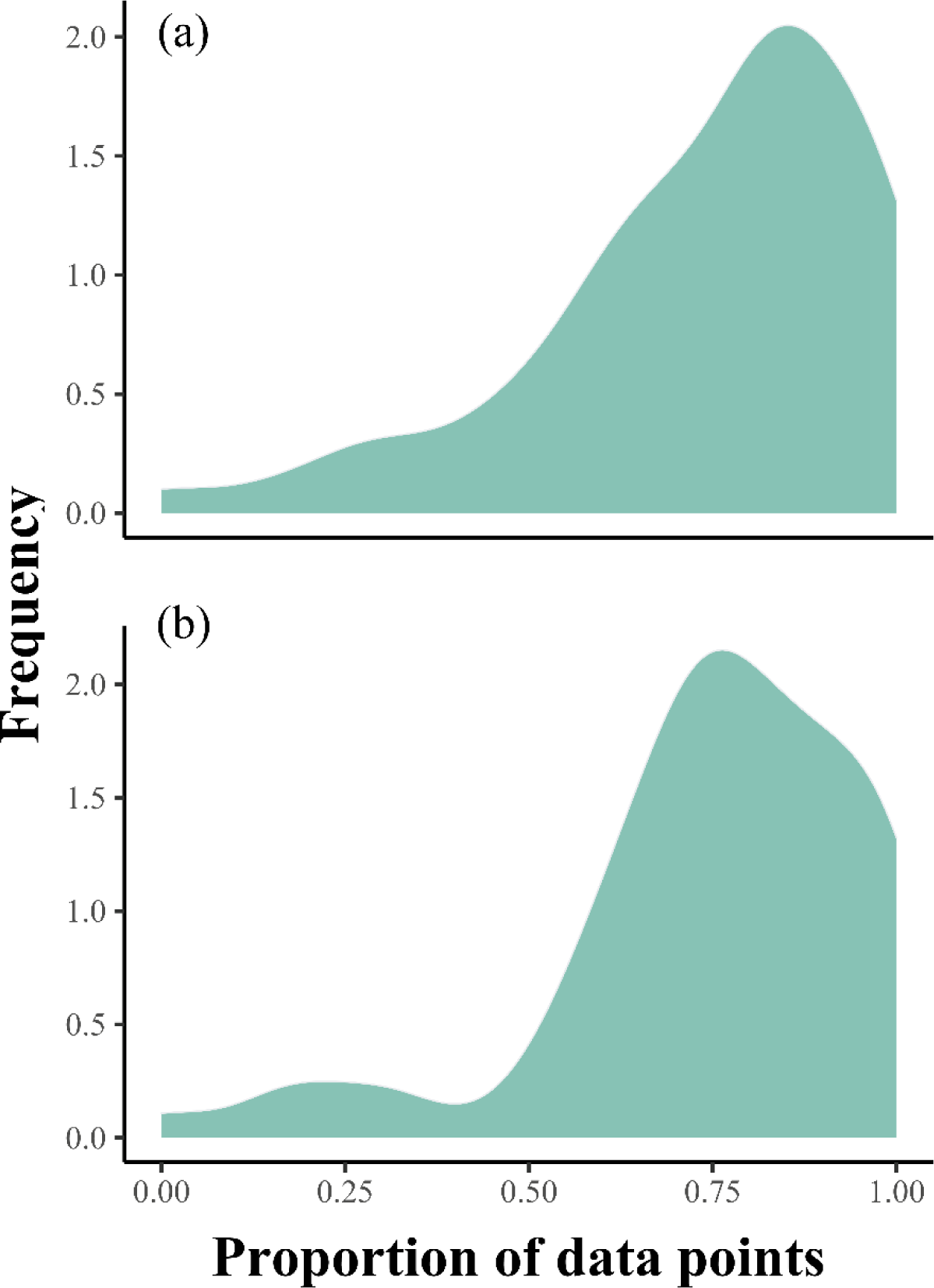
Hind-casting distributions of annual herbs before 1950. Density plot showed the proportions of actual species distribution points that fell within the predicted ranges before 1950 by abiotic SDM (a) and phenology-informed SDMs (b), respectively.

**Fig. S3.**
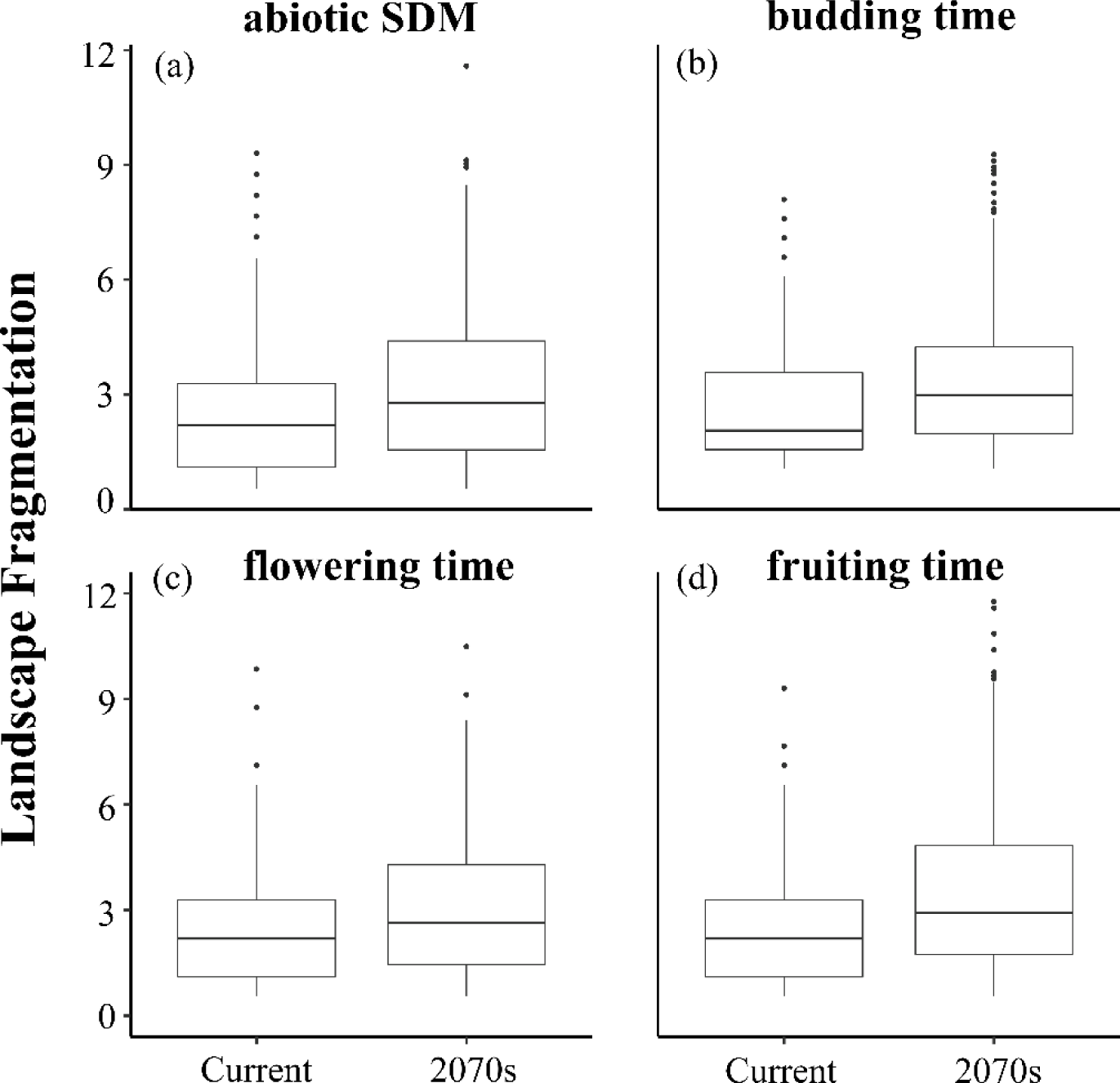
Landscape fragmentation index of species’ current and future distribution ranges. Species distributions were separately estimated based on the abiotic species distribution models (SDMs) (a) and phenology-informed SDMs (b-d). We quantified plant phenology using three indices: the day of year for peak budding (b), peak flowering (c) and peak fruiting (d).

**Fig. S4.**
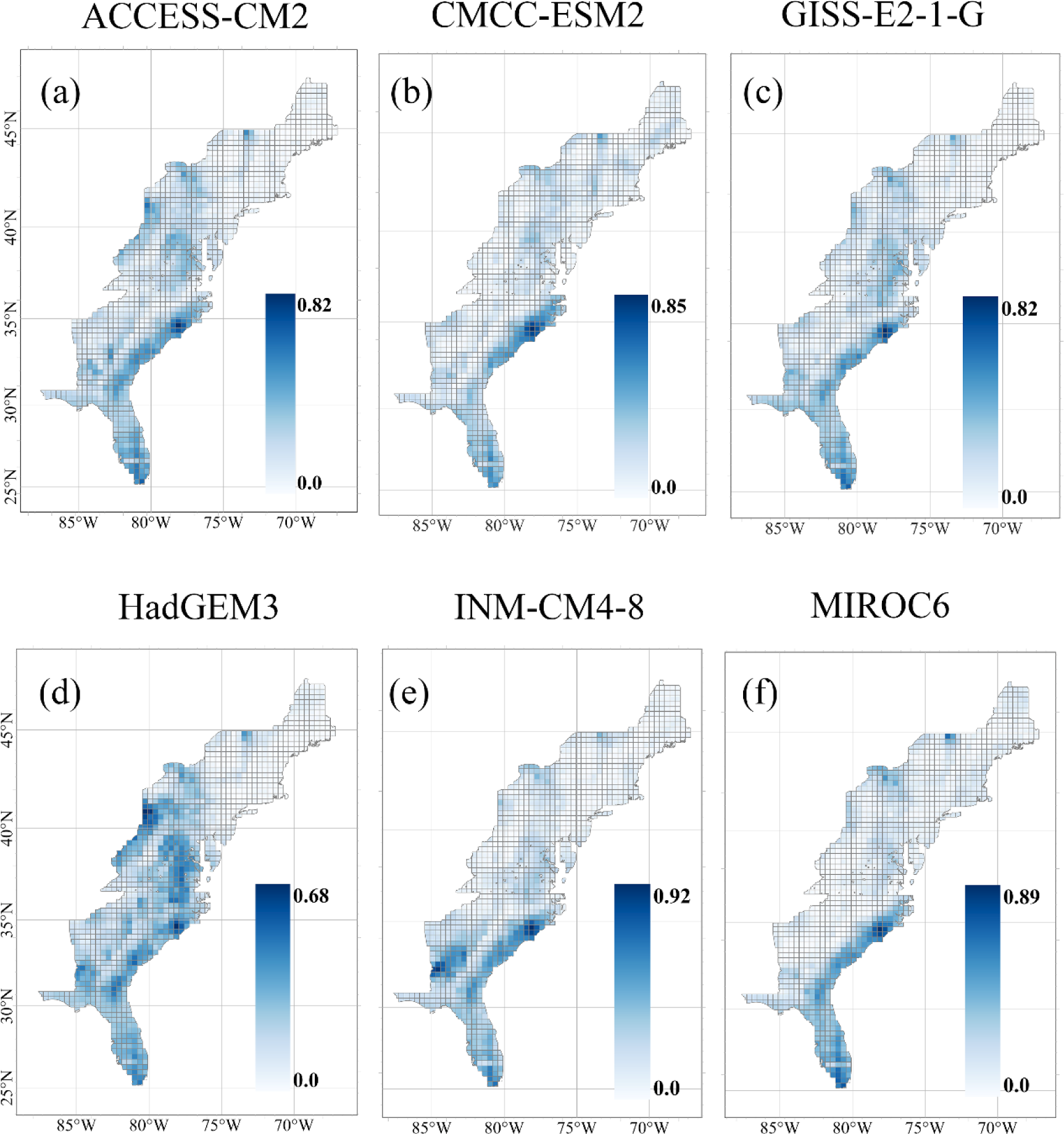
Geographical patterns in the proportion of species gains between current and forecasted species distributions in the 2070s under different General Circulation Models (GCMs). Species gain refers to regions (i.e., grid cells) where species are predicted to be absent currently but are forecasted to be present in the 2070s.

**Fig. S5.**
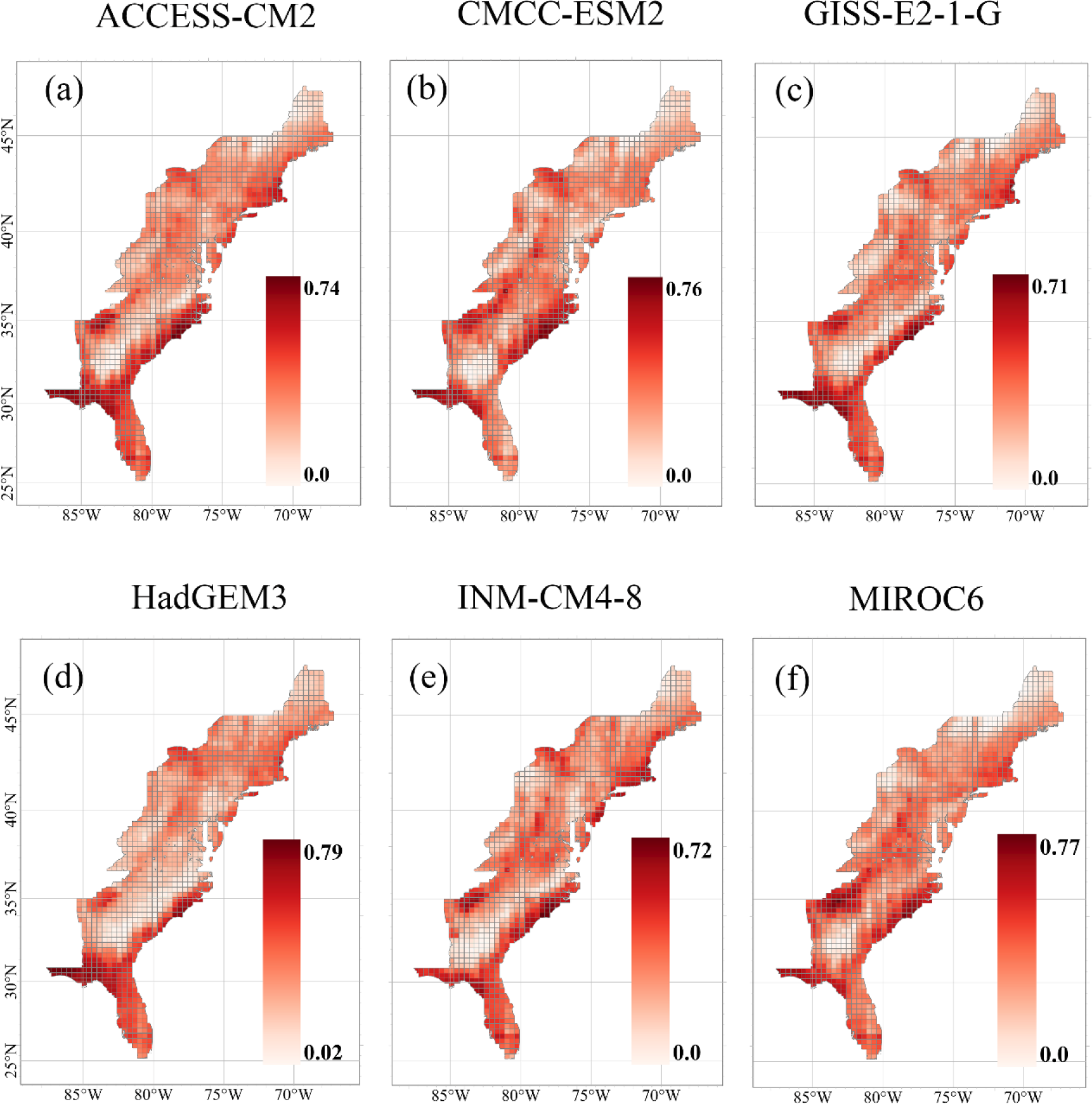
Geographical patterns in the proportion of species losses between current and forecasted species distributions in the 2070s under different General Circulation Models (GCMs). Species loss refers to regions (i.e., grid cells) where species are predicted to be present currently but are forecasted to be absent in the 2070s.

**Fig. S6.**
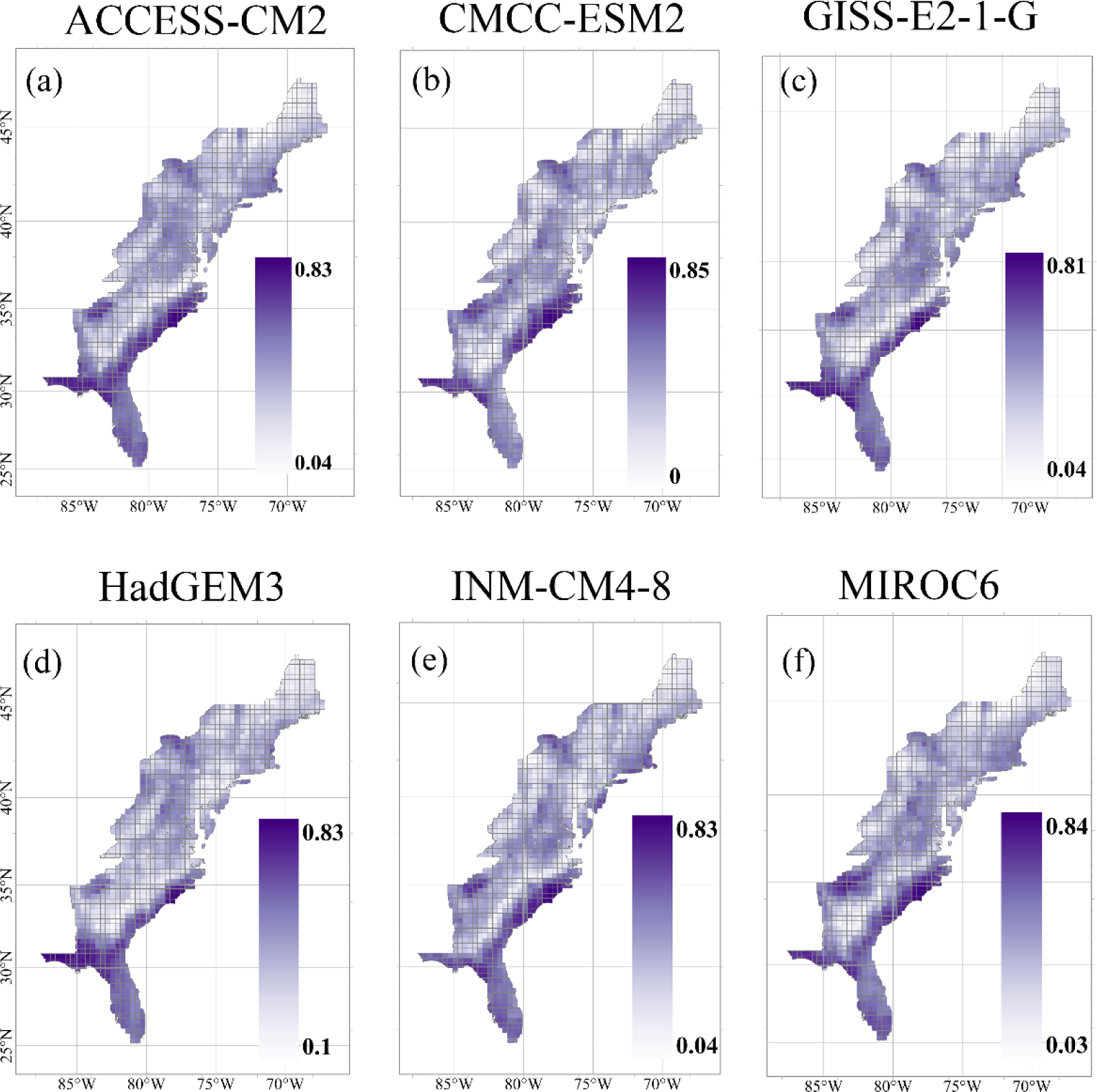
Geographical patterns in the proportion of species turnover between current and forecasted species distributions in the 2070s under different General Circulation Models (GCMs). Species turnover is simply defined as the relative changes in species composition within each grid cell.

**Fig. S7.**
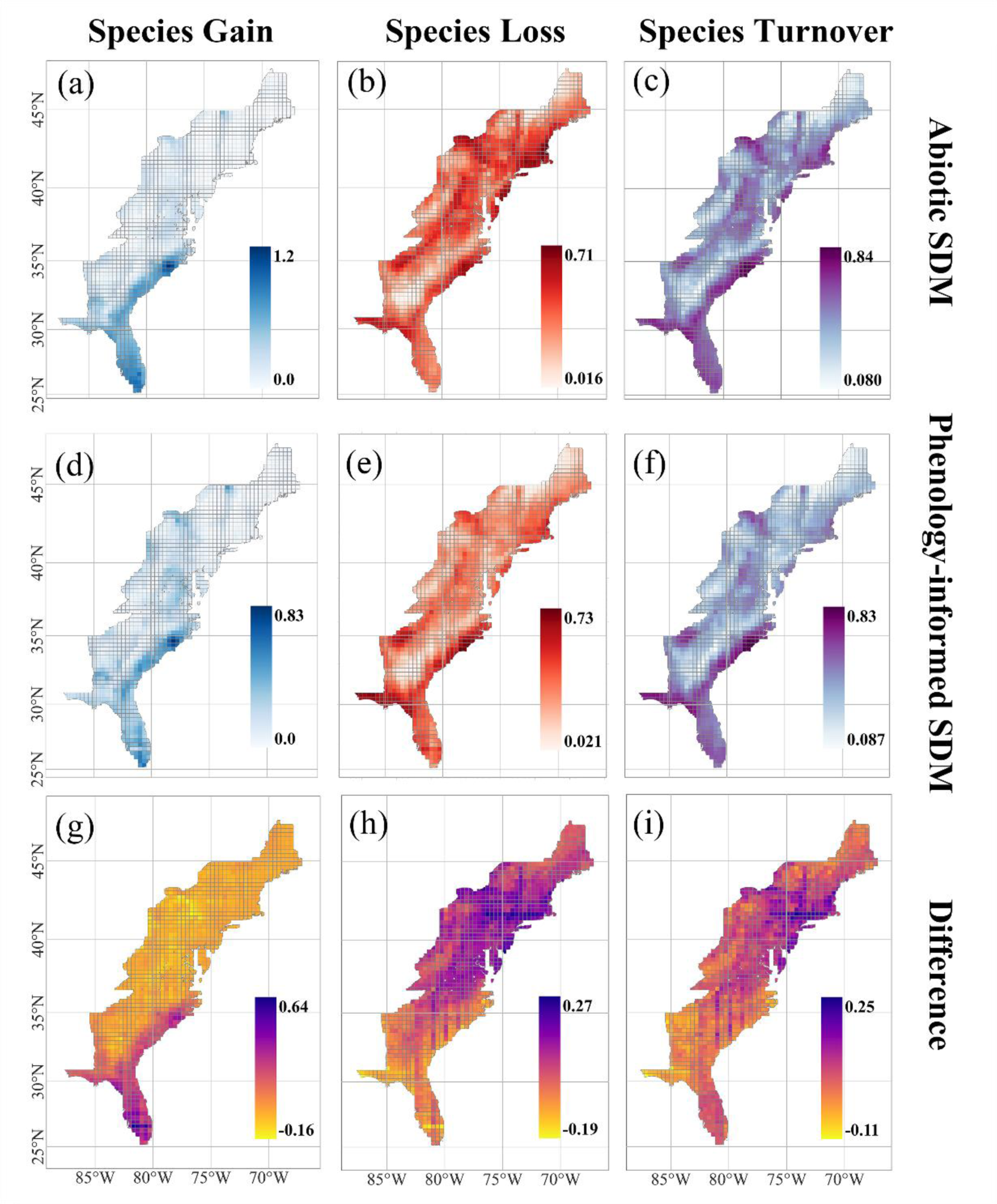
Geographical patterns in the proportion of species gains, losses and turnover between current and forecasted species distributions in the 2070s based on the abiotic SDMs (a-c) and phenology-informed SDMs (d-f), respectively. The differences in the proportion of species gains, losses and turnover between the abiotic SDM and phenology-informed SDMs are also shown (g-i).

**Fig. S8.**
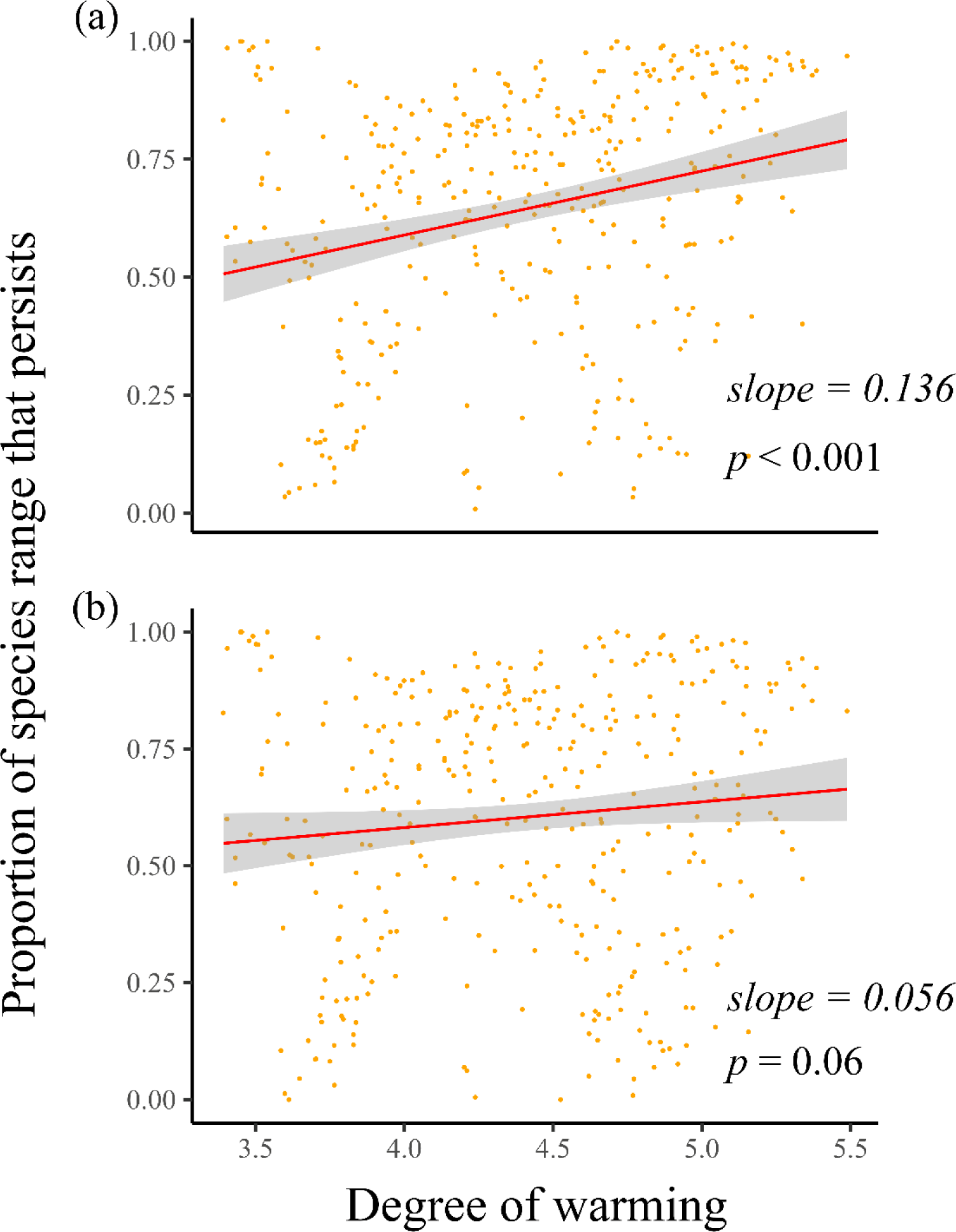
Relationships between the mean degree of warming predicted across the range of species and the proportion of species’ current geographic range that were projected to persist under future climate change scenarios. Species future geographic distributions were forecasted by phenology-informed (a) and abiotic SDMs (b), respectively.

**Fig. S9.**
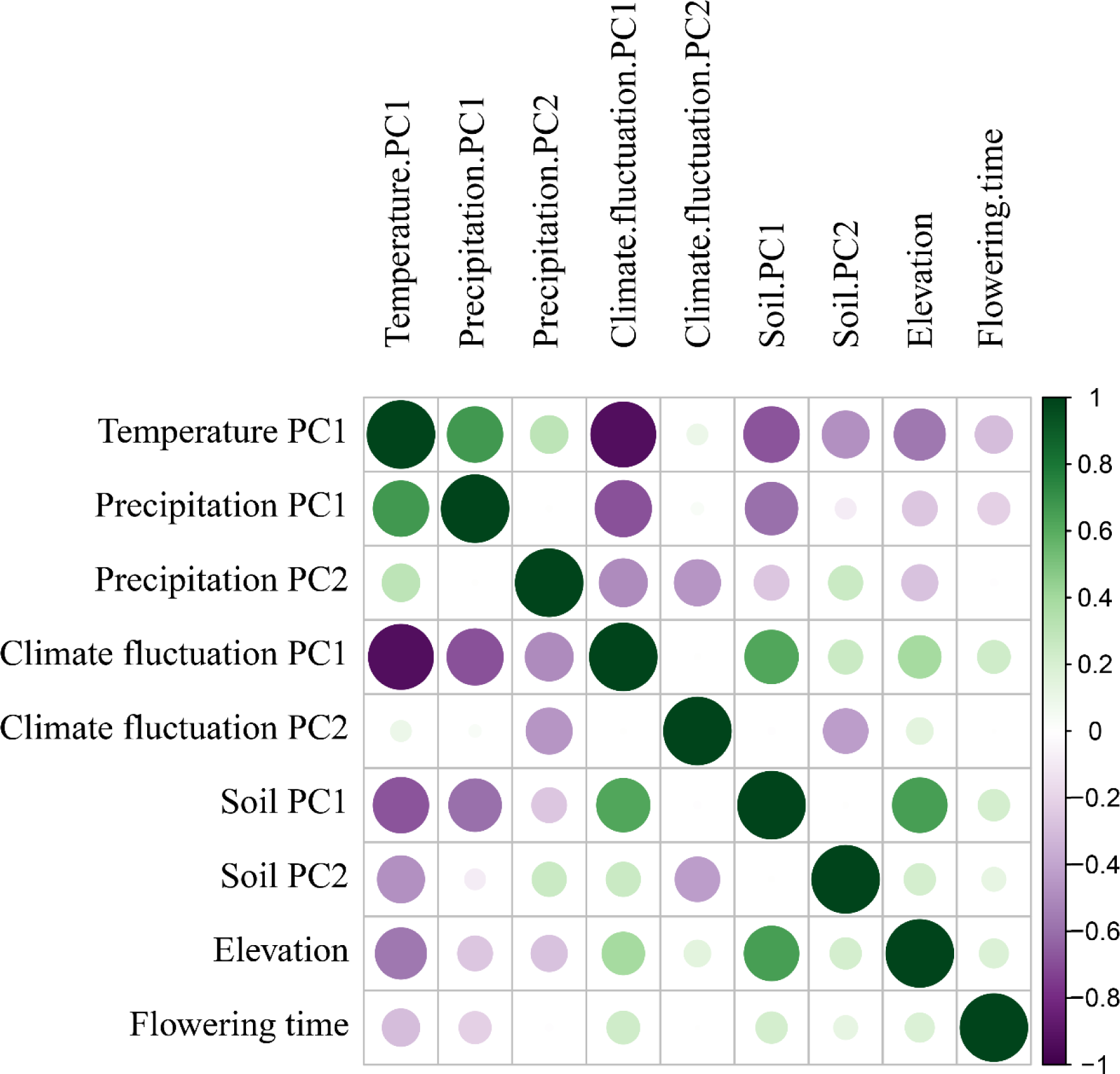
Correlations among environmental variables and phenology predictors used for building phenology-informed species distribution models. We used seven principal component scores as predictor variables representing mean temperature, mean precipitation, climatic fluctuation and soils. Flowering time refers to the day of year (DOY) for plant peak flowering.

**Table S1.** Summary of linear mixed models for plant budding, flowering and fruiting. Marginal and Conditional *R*^2^ values for each model are also provided. The “:” symbol in the term column represents an interaction between two variables.

| Term | Budding | Flowering | Fruiting |
| --- | --- | --- | --- |
| Intercept | 220.75 ± 12.64 *** | 226.10 ± 10.20 *** | 187.94 ± 12.85 *** |
| bio1 | -7.84 ± 3.57 *** | -4.532 ± 3.50 | -5.01 ± 3.89 |
| bio4 | 5.86 ± 2.58 *** | 12.59 ± 2.42*** | 11.78 ± 2.98 *** |
| bio12 | 1.58 ± 1.01 | 0.85 ± 1.01 | 0.28 ± 1.20 |
| bio15 | -0.24 ± 1.01 | 1.65 ± 1.01 | 3.00 ± 1.27 *** |
| Year | -0.004 ± 0.005 | -0.012 ± 0.003*** | 0.019 ± 0.005 *** |
| Herbaceous perennial | -29.73 ± 6.91 *** | -35.27 ± 7.15 *** | -27.16 ± 6.79 *** |
| Vine perennial | -40.37 ± 14.20 *** | -40.56 ± 14.70 *** | -25.38 ± 14.27 |
| Woody | -37.20 ± 8.21 *** | -41.57 ± 8.51 *** | -16.82 ± 8.03 *** |
| Native | -10.64 ± 7.38 | 0.87 ± 7.66 | -2.89 ± 7.39 |
| bio1: Herbaceous perennial | -9.89 ± 3.16 *** | -4.99 ± 3.01 | -9.41 ± 3.38 *** |
| bio1: Vine perennial | -7.13 ± 6.48 | 2.91 ± 6.06 | -7.14 ± 6.91 |
| bio1: Woody | -10.03 ± 3.87 *** | -8.77 ± 3.67 *** | -11.94 ± 3.98 *** |
| bio4: Herbaceous perennial | -3.74 ± 2.27 | -5.07 ± 2.01 *** | -5.95 ± 2.56 *** |
| bio4: Vine perennial | -6.24 ± 4.57 | -0.1 ± 4.00 | -10.67 ± 5.25 *** |
| bio4: Woody | 1.65 ± 2.77 | -4.08 ± 2.43 | -5.74 ± 2.99 |
| bio12: Herbaceous perennial | 0.03 ± 0.89 | -1.51 ± 0.84 | 0.38 ± 1.02 |
| bio12: Vine perennial | 0.35 ± 1.78 | -1.07 ± 1.69 | 1.23 ± 2.11 |
| bio12: Woody | 0.06 ± 1.09 | -1.07 ± 1.02 | -0.62 ± 1.19 |
| bio15: Herbaceous perennial | 0.76 ± 0.89 | -1.46 ± 0.84 | 0.01 ± 1.09 |
| bio15: Vine perennial | 2.83 ± 1.81 | 0.78 ± 1.68 | 0.55 ± 2.23 |
| bio15: Woody | -0.34 ± 1.10 | -2.97 ± 1.02 *** | -1.39 ± 1.26 |
| bio1: Native | -0.14 ± 3.49 | -6.07 ± 3.37 | -1.10 ± 3.90 |
| bio4: Native | -2.39 ± 2.50 | -6.99 ± 2.30 *** | -3.66 ± 2.97 |
| bio12: Native | -1.76 ± 0.98 | 0.25 ± 0.96 | -0.17 ± 1.20 |
| bio15: Native | 0.22 ± 0.99 | 0.22 ± 0.97 | -2.58 ± 1.27 *** |
| Marginal $R^2$ | 0.247 | 0.227 | 0.142 |
| Conditional $R^2$ | 0.423 | 0.521 | 0.428 |

**Table S2.** Environmental variables used in conducting principal component analyses (PCA) integrated into phenology-informed species distribution models (SDMs)

| **Bioclimatic Variable** | **Full Name** | **Categories** |
| --- | --- | --- |
| bio1 | Annual Mean Temperature | Mean Temperature |
| bio2 | Mean Diurnal Range | Climatic Fluctuation |
| bio3 | Isothermality | Climatic Fluctuation |
| bio4 | Temperature Seasonality | Climatic Fluctuation |
| bio5 | Max Temperature of Warmest Month | Mean Temperature |
| bio6 | Min Temperature of Coldest Month | Mean Temperature |
| bio7 | Temperature Annual Range | Climatic Fluctuation |
| bio8 | Mean Temperature of Wettest Quarter | Mean Temperature |
| bio9 | Mean Temperature of Driest Quarter | Mean Temperature |
| bio10 | Mean Temperature of Warmest Quarter | Mean Temperature |
| bio11 | Mean Temperature of Coldest Quarter | Mean Temperature |
| bio12 | Annual Precipitation | Mean Precipitation |
| bio13 | Precipitation of Wettest Month | Mean Precipitation |
| bio14 | Precipitation of Driest Month | Mean Precipitation |
| bio15 | Precipitation Seasonality | Climatic Fluctuation |
| bio16 | Precipitation of Wettest Quarter | Mean Precipitation |
| bio17 | Precipitation of Driest Quarter | Mean Precipitation |
| bio18 | Precipitation of Wettest Quarter | Mean Precipitation |
| bio19 | Precipitation of Coldest Quarter | Mean Precipitation |
| SC | Sand Content | Soil |
| CC | Clay Content | Soil |
| SC | Silt Content | Soil |
| BD | Bulk Density | Soil |
| CFR | Coarse Fragments | Soil |

**Table S3.** Results of tests for phylogenetic signal on predicted changes in species’ habitat suitability. Range persistence: current occupied range predicted to have high future suitability; range retreat: current occupied range predicted to have low future suitability; range expansion: current unoccupied range predicted to have high future suitability. Phylogenetic signals are examined by randomization tests for Blomberg’s *K* and likelihood ratio tests for Pagel’s lambda.

| Phenological traits |  | Blomberg's *K* |  | Pagel's lambda |  |
| --- | --- | --- | --- | --- | --- |
|  |  | Value | *p*-value | Value | *p*-value |
| budding | range persistence | < 0.001 | 0.879 | < 0.001 | 1 |
|  | range retreat | < 0.001 | 0.879 | < 0.001 | 1 |
|  | range expansion | 0.004 | 0.367 | 0.258 | 0.001* |
| flowering | range persistence | < 0.001 | 0.928 | 0.023 | 1 |
|  | range retreat | < 0.001 | 0.915 | 0.023 | 1 |
|  | range expansion | 0.003 | 0.284 | 0.219 | 0.002* |
| fruiting | range persistence | < 0.001 | 0.857 | < 0.001 | 1 |
|  | range retreat | < 0.001 | 0.854 | < 0.001 | 1 |
|  | range expansion | 0.001 | 0.541 | 0.207 | 0.002* |

**Table S4.**
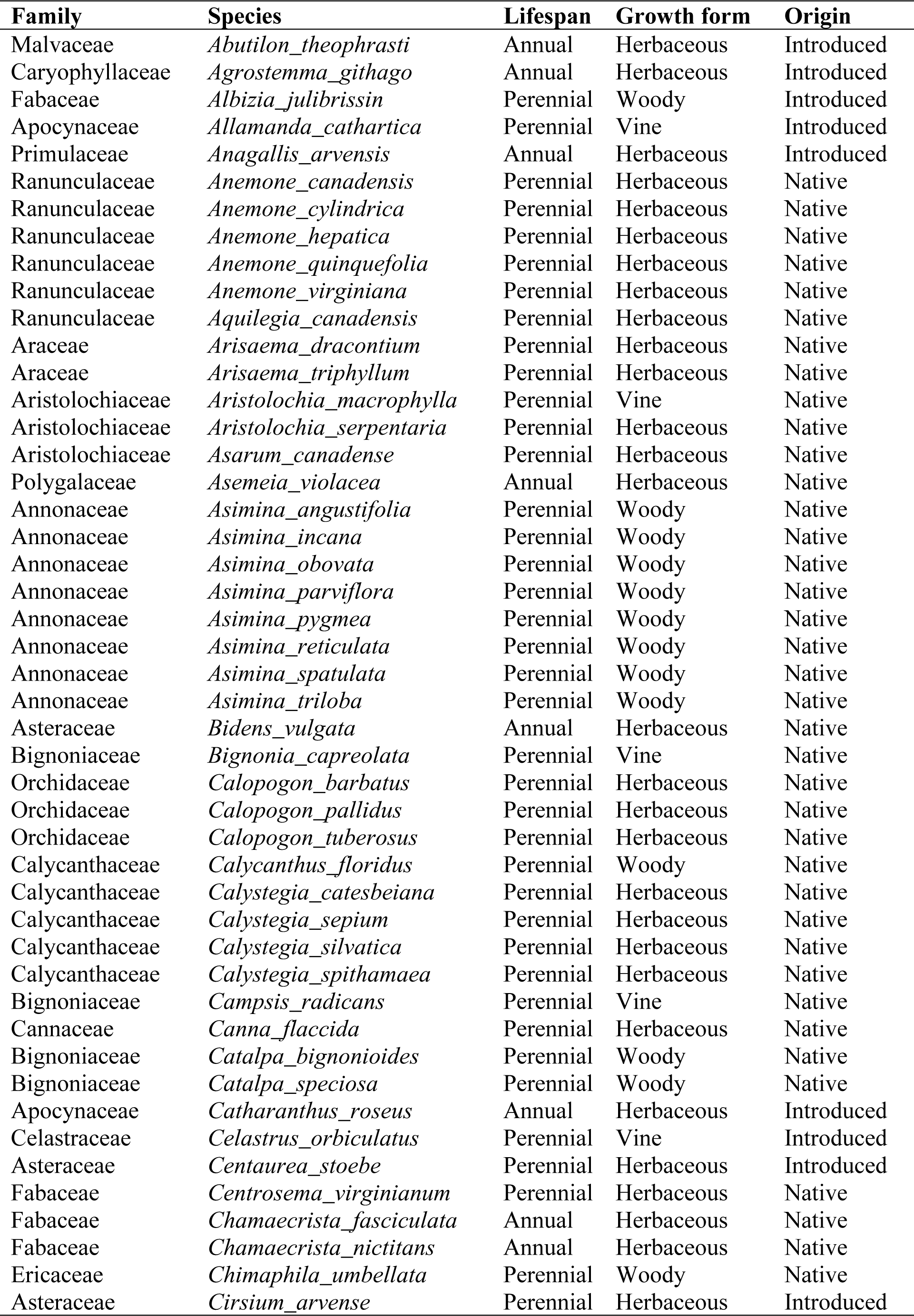
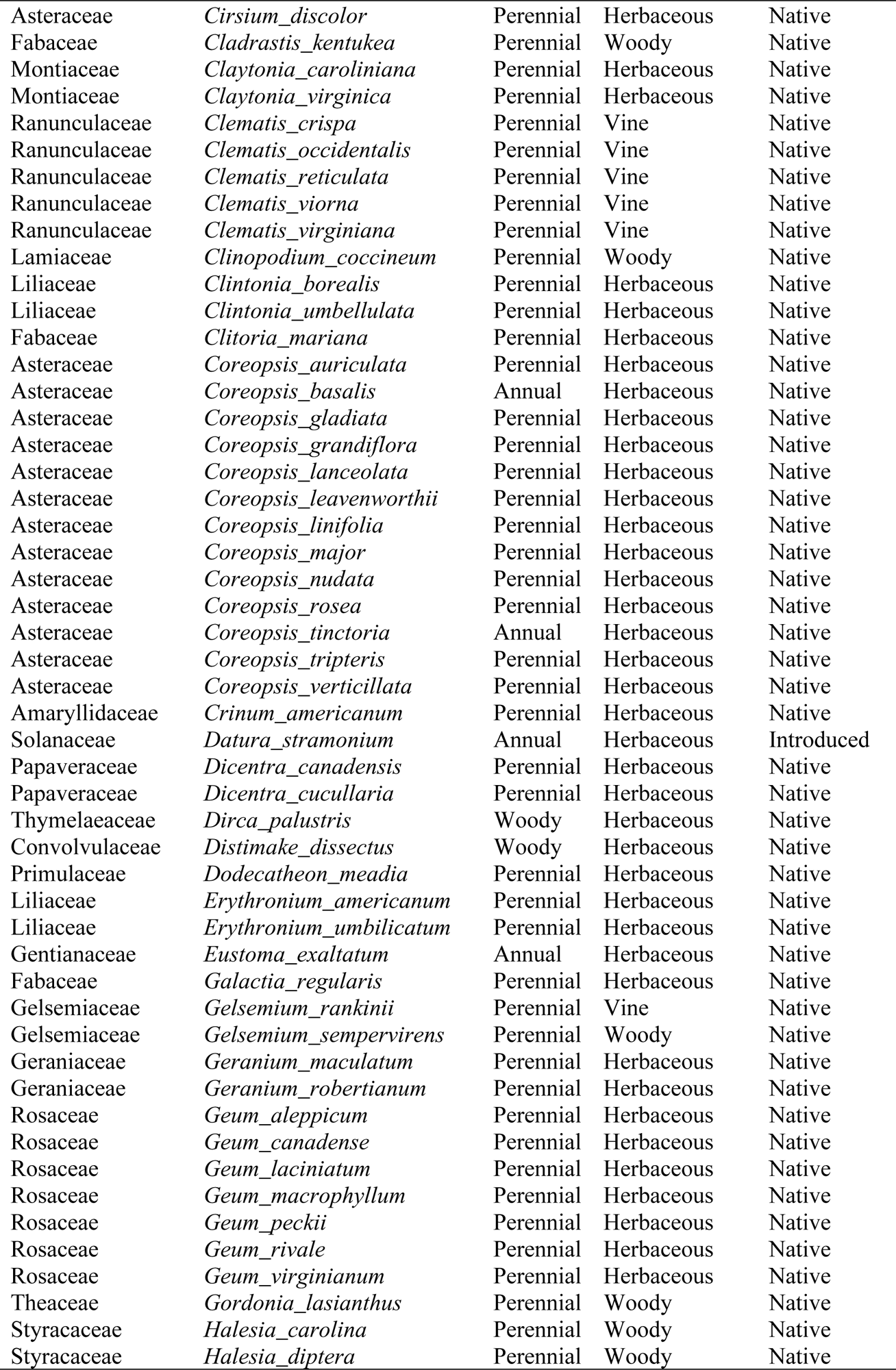
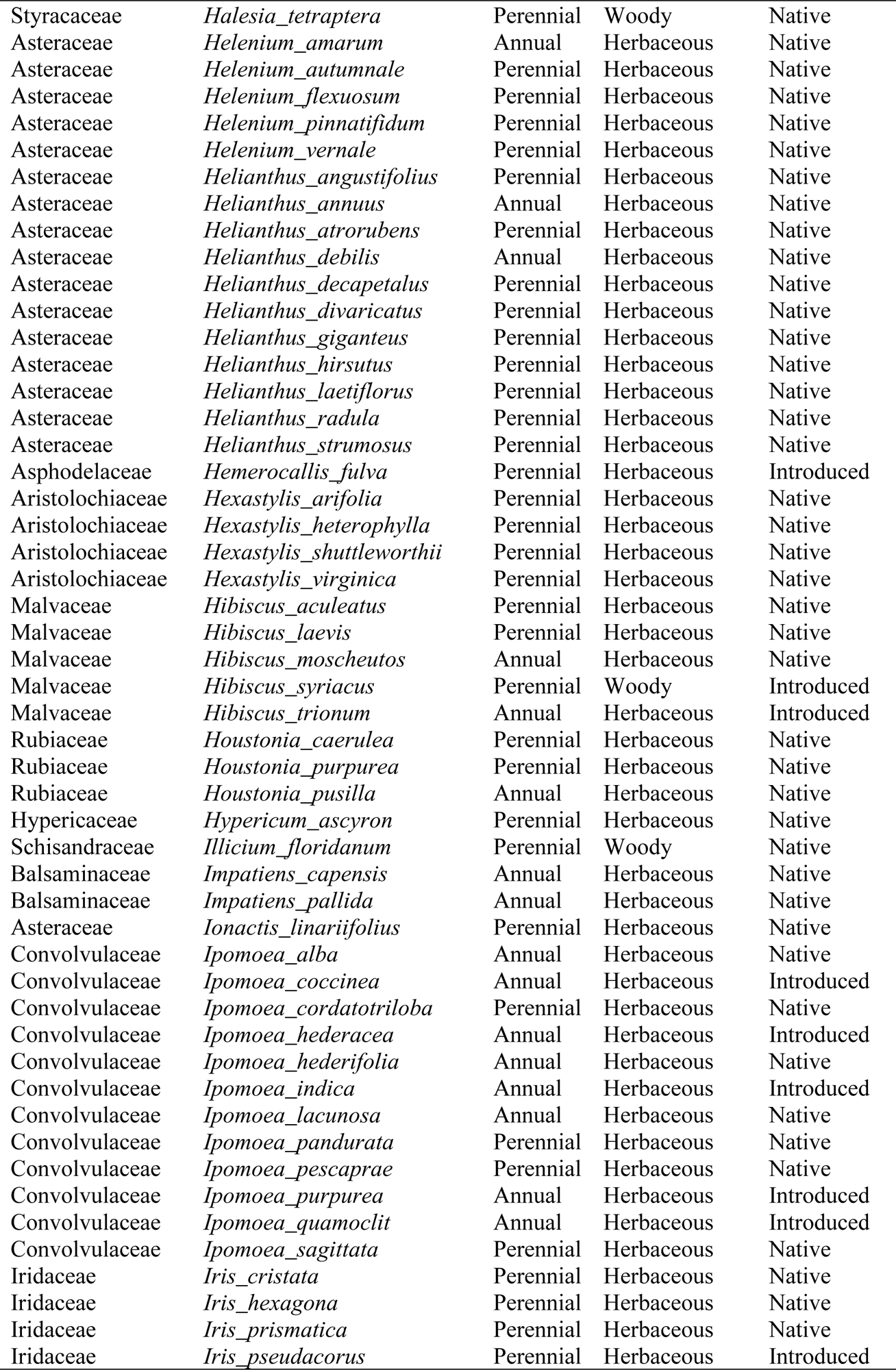
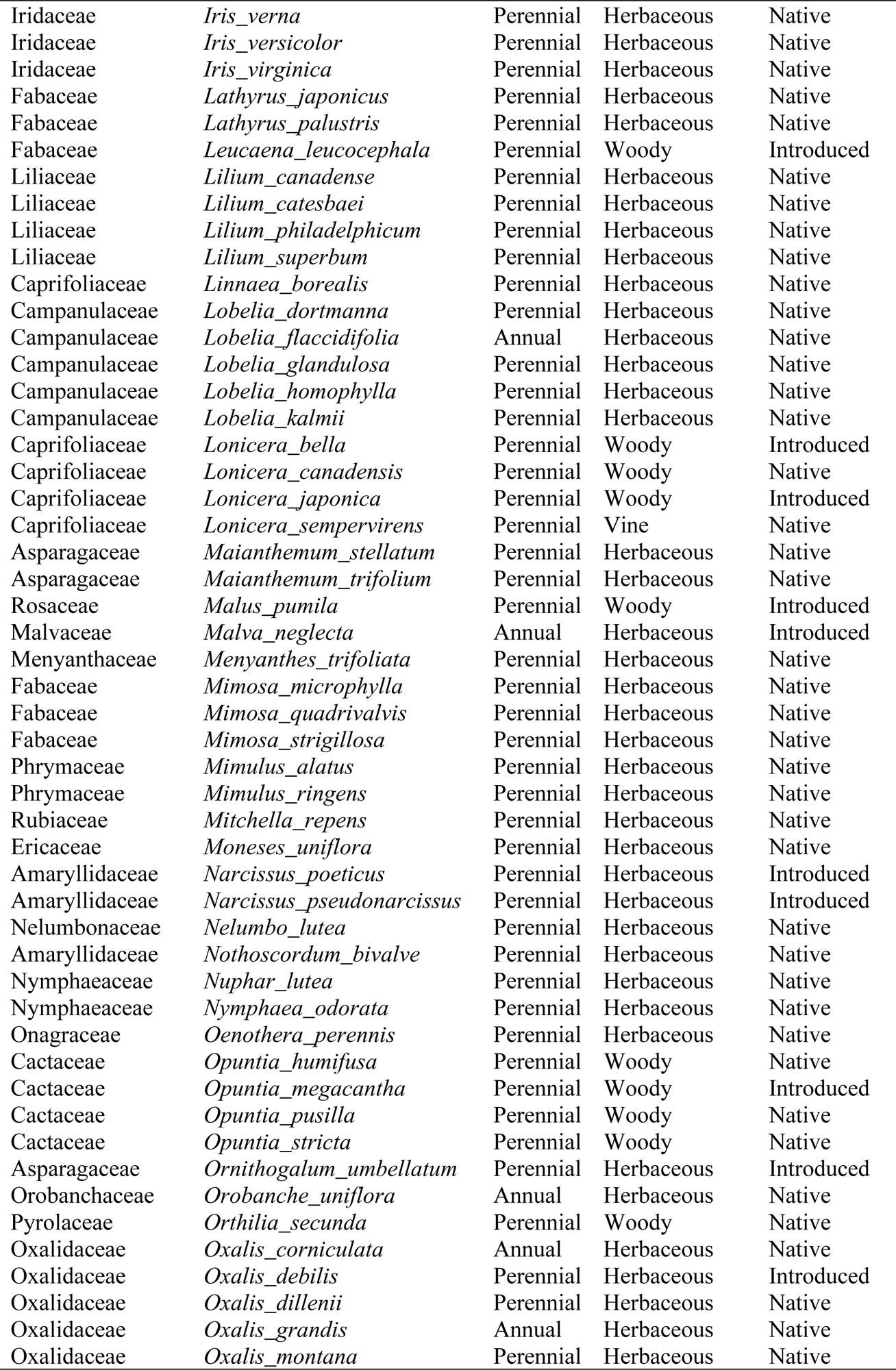
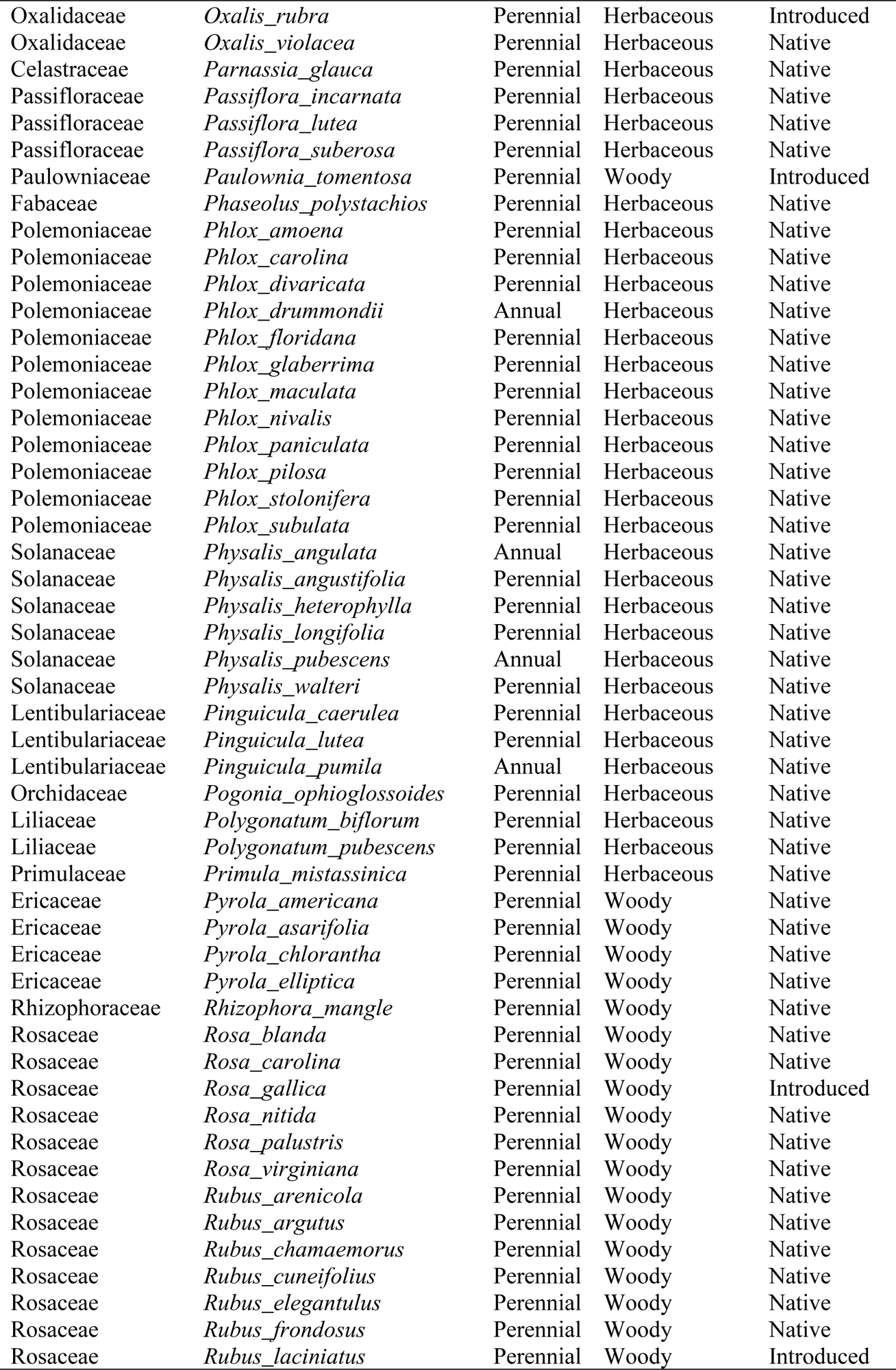
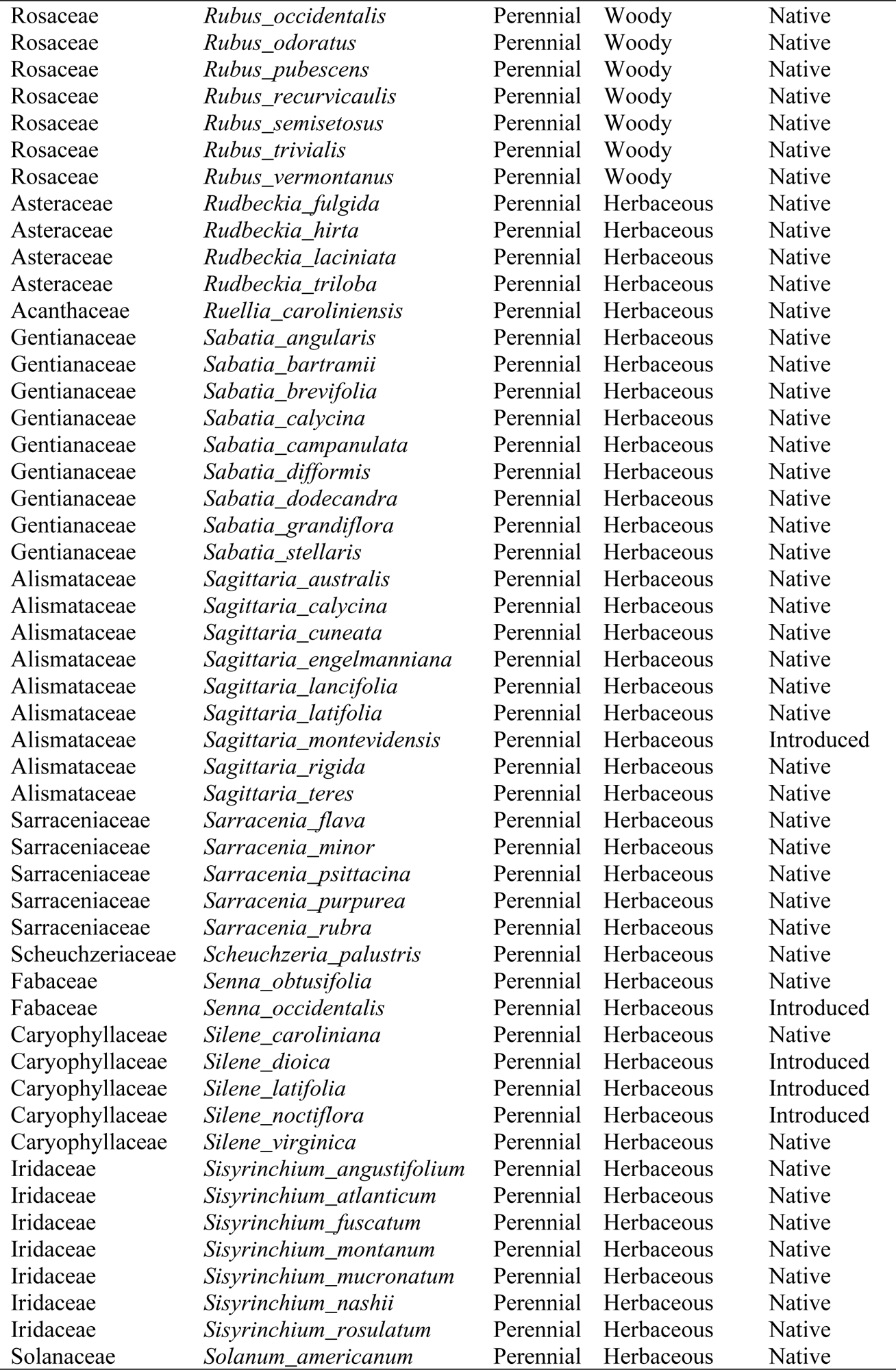
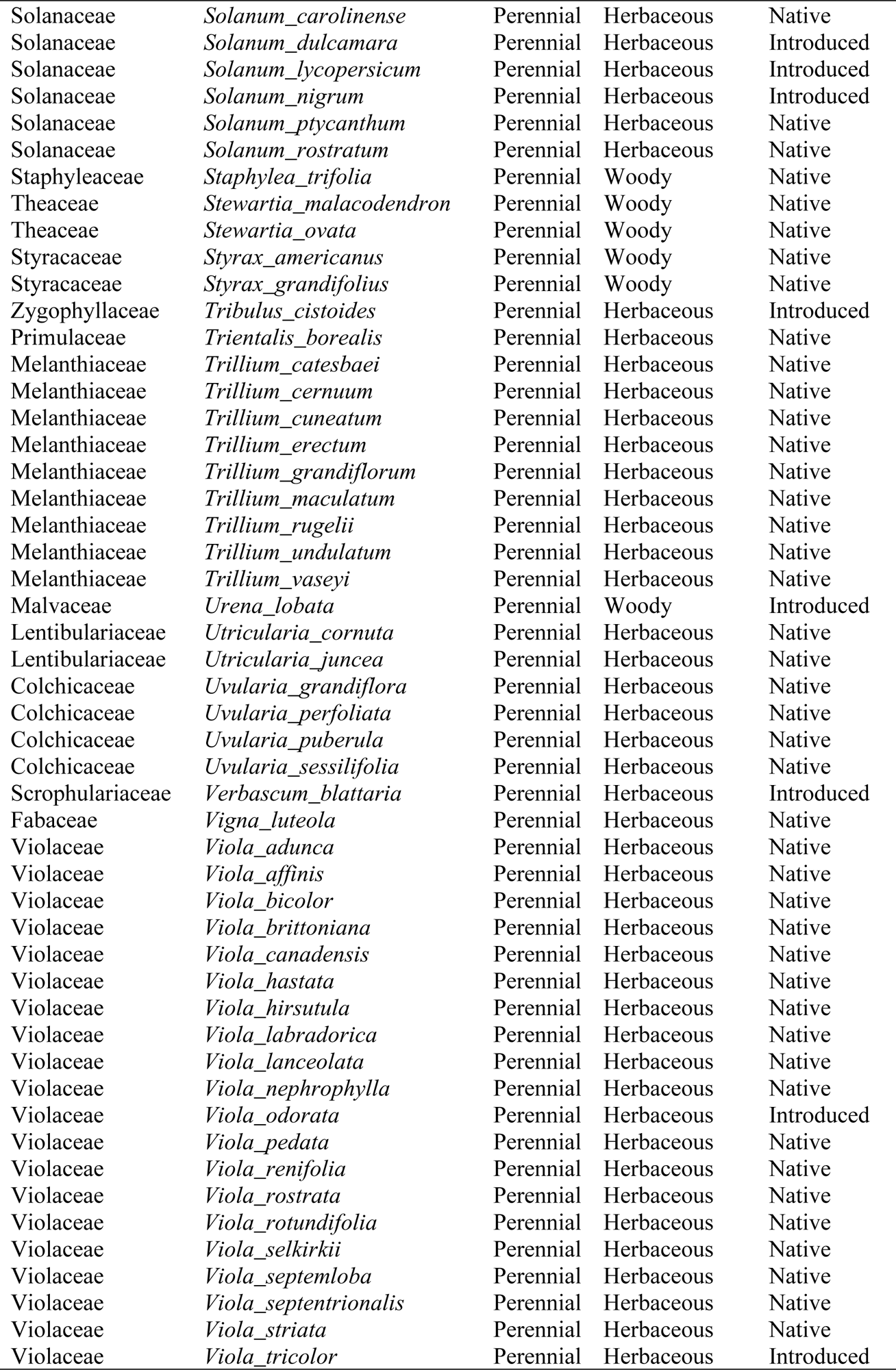
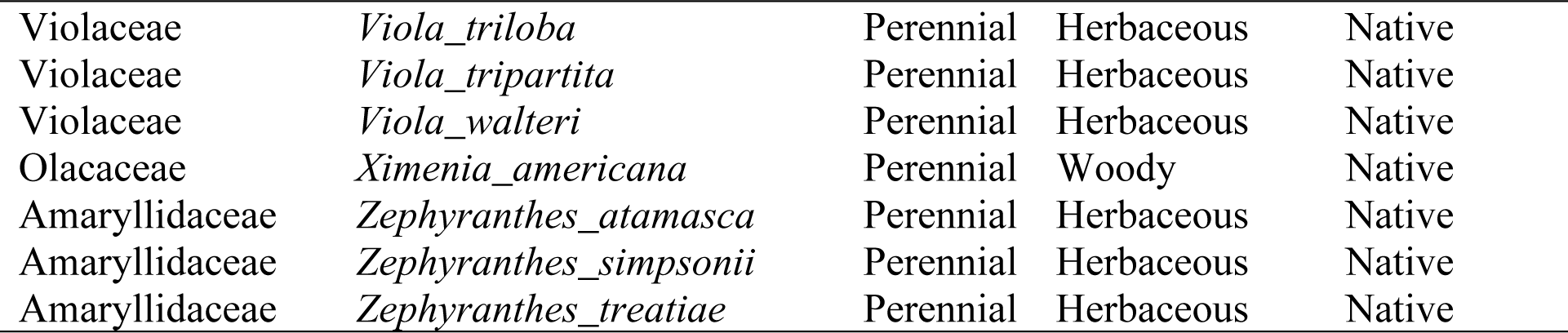
Taxonomic, lifespan, growth form, and origin information for the 360 plant species used in our study

**Table S5.** Phylogenetic signals of the mean peak budding, flowering and fruiting time and the responses of peak budding, flowering and fruiting time to environmental variables.

| Phenological traits | Blomberg's K |  | Pagel's lambda |  |
| --- | --- | --- | --- | --- |
|  | Value | *P* | Value | *P* |
| **Peak budding time** |  |  |  |  |
| Phenological sensitivity to bio1 | 0.001 | 0.94 | 0.168 | 0.004 |
| Phenological sensitivity to bio4 | 0.008 | 0.015 | 0.09 | 0.259 |
| Phenological sensitivity to bio12 | 0.003 | 0.347 | < 0.001 | 1 |
| Phenological sensitivity to bio15 | 0.002 | 0.595 | < 0.0001 | 1 |
| Median budding time | 0.007 | 0.014 | 0.825 | < 0.001 |
| **Peak flowering time** |  |  |  |  |
| Phenological sensitivity to bio1 | 0.001 | 0.908 | 0.247 | < 0.001 |
| Phenological sensitivity to bio4 | 0.003 | 0.301 | < 0.001 | 1 |
| Phenological sensitivity to bio12 | 0.006 | 0.055 | 0.554 | 0.173 |
| Phenological sensitivity to bio15 | 0.001 | 0.924 | < 0.001 | 1 |
| Median flowering time | 0.015 | 0.001 | 0.87 | < 0.001 |
| **Peak fruiting time** |  |  |  |  |
| Phenological sensitivity to bio1 | 0.004 | 0.171 | 0.447 | < 0.001 |
| Phenological sensitivity to bio4 | 0.005 | 0.036 | 0.096 | 0.089 |
| Phenological sensitivity to bio12 | 0.001 | 0.745 | 0.081 | 0.155 |
| Phenological sensitivity to bio15 | 0.002 | 0.465 | < 0.001 | 1 |
| Median fruiting time | 0.0071 | 0.008 | 0.821 | < 0.001 |

**Table S6.**
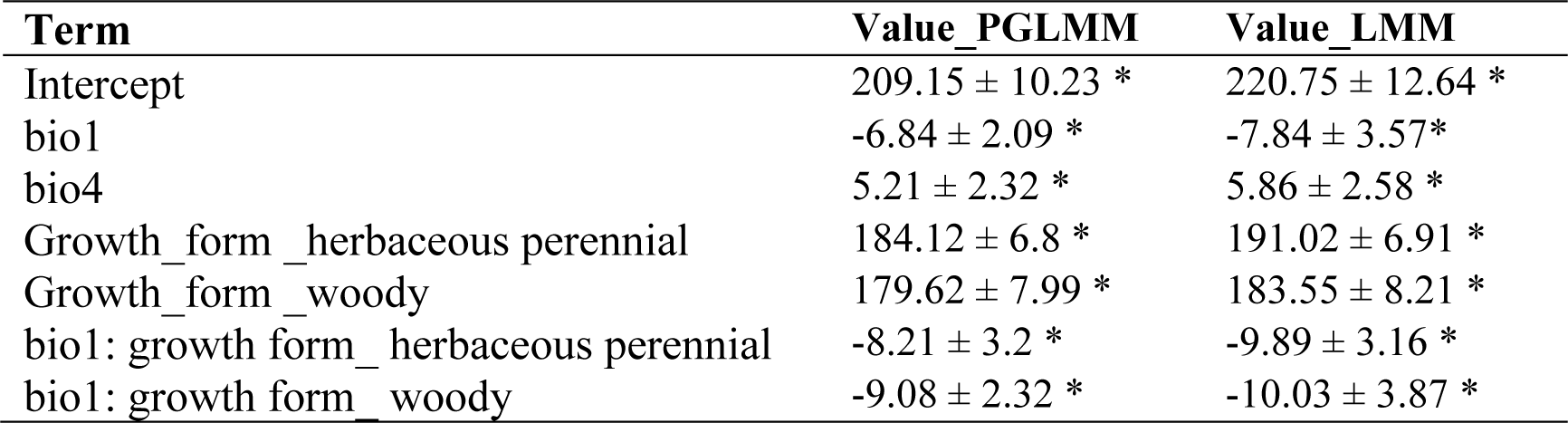
Summary of phylogenetic linear mixed models for plant peak budding time.

**Table S7.** Summary of phylogenetic linear mixed models for plant peak flowering time.

| **Term** | **Value_PGLMM** | **Value_LMM** |
| --- | --- | --- |
| Intercept | 224.50 ± 12.9 * | 226.1 ± 10.20 * |
| bio4 | 8.77 ± 2.32 * | 12.59 ± 2.42 * |
| Growth_form _herbaceous perennial | 193.24 ± 8.2 * | 190.83 ± 7.15 * |
| Growth_form _woody | 189.28 ± 12.9 * | 184.53 ± 8.51 * |
| bio1: growth form_ woody | -9.00 ± 3.2 * | -8.77 ± 3.67 * |
| bio4: growth form_ herbaceous perennial | -6.01 ± 2.7 * | -5.07 ± 2.01 * |
| bio15: growth form_ woody | -1.84 ± 0.68 * | -2.97 ± 1.02 * |
| bio4: status_ native | -6.72 ± 2.89 * | -6.99 ± 2.30 * |

**Table S8.** Summary of phylogenetic linear mixed models for plant peak fruiting time.

| **Term** | **Value_PGLMM** | **Value_LMM** |
| --- | --- | --- |
| Intercept | 193.53 ± 10.01 * | 187.94 ± 12.85 * |
| bio4 | 9.00 ± 2.67 * | 11.78 ± 2.98 * |
| bio15 | 2.79 ± 1.07 * | 3.00 ± 1.27 * |
| Growth_form _herbaceous perennial | 158.16 ± 6.20 * | 160.78 ± 6.79 * |
| Growth_form _woody | 170.33 ± 5.62 * | 171.12 ± 8.03 * |
| bio1: growth form_herbaceous perennial | -12.23 ± 5.66 * | -9.41 ± 3.38 * |
| bio1: growth form_ woody | -9.62 ± 3.66 * | -11.94 ± 3.98 * |
| bio4: growth form_herbaceous perennial | -5.20 ± 1.46 * | -5.95 ± 2.56 * |
| bio15: status_ native | -2.02 ± 0.89* | -2.58 ± 1.27 * |

## Notes

### Competing Interest Statement

The authors have declared no competing interest.

## References

1. Gottfried, M., Pauli, H., Reiter, K. & Grabherr, G. Potential effects of climate change on alpine and nival plants in the Alps. In: Mountain Biovidersity-A Global Assessment (eds. C Körner, EM Spehn), pp.213–223. Parthenon Publishing, London, New York.

2. Kelly, A. E. & Goulden, M. L. Rapid shifts in plant distribution with recent climate change. Proc. Natl Acad. Sci. 105, 11823–11826 (2008).

3. Parker, J. Cold resistance in woody plants. The Botanical Review 29, 123–201(1963).

4. Whittaker, R. H. Communities and ecosystems. (Macmillan, 1970).

5. Kotta, J. et al. Integrating experimental and distribution data to predict future species patterns. Sci. Rep. 9, 1–14 (2019).

6. Mi, C. et al. Global Protected Areas as refuges for amphibians and reptiles under climate change. Nat. Commun. 14, 1389 (2023).

7. Thomas, C. D. et al. Extinction risk from climate change. Nature 427, 145–148 (2004).

8. Hutchinson, G. E. Concluding remarks. Cold spring harbor symposia on quantitative biology. 22, 415–427(1957).

9. Benito Garzön, M., Robson, T. M. & Hampe, A. ΔTraitSDMs: species distribution models that account for local adaptation and phenotypic plasticity. New Phytol. 222, 1757–1765 (2019).

10. Kearney, M. & Porter, W. Mechanistic niche modelling: combining physiological and spatial data to predict species’ ranges. Ecol. Lett. 12, 334–350 (2009).

11. Rosenzweig, M. L. Habitat selection as a source of biological diversity. Evol. Biol. 1, 315–330 (1987).

12. Westoby, M. & Wright, I. J. Land-plant ecology on the basis of functional traits. Trends Ecol. Evol. 21, 261–268 (2006).

13. Carboni, M. et al. Functional traits modulate the response of alien plants along abiotic and biotic gradients. Glob. Ecol. Biogrogr. 27, 1173–1185 (2018).

14. Pollock, L. J., Morris, W. K. & Vesk, P. A. The role of functional traits in species distributions revealed through a hierarchical model. Ecography 35, 716–725 (2012).

15. Evans, M. E. K., Merow, C., Record, S., McMahon, S. M. & Enquist, B. J. Towards Process-based Range Modeling of Many Species. Trends Ecol. Evol. 31, 860–871 (2016).

16. Chuine, I. & Beaubien, E. G. Phenology is a major determinant of tree species range. Ecol. Lett. 4, 500–510 (2001).

17. Meineke, E. K., Davis, C. C. & Davies, T. J. Phenological sensitivity to temperature mediates herbivory. Glob. Change Biol. 27, 2315–2327 (2021).

18. Schwartz, M. D. (ed.) Phenology: an integrative environmental science. (Kluwer Academic, 2003).

19. O’Neil, P. & Schmitt, J. Natural selection on genetically correlated phenological characters in *Lythrum salicaria*. Evolution 47, 267–274 (1997).

20. Chuine, I. Why does phenology drive species distribution? Phil. Trans. R. Soc. B 365, 3149–3160 (2010).

21. Hereford, J., Schmitt, J. & Ackerly, D. D. The seasonal climate niche predicts phenology and distribution of an ephemeral annual plant, *Mollugo verticillata*. J. Ecol. 105, 1323–1334 (2017).

22. Willis, C. G., Ruhfel, B., Primack, R. B., Miller-Rushing, A. J. & Davis, C. C. Phylogenetic patterns of species loss in Thoreau’s woods are driven by climate change. Proc. Natl Acad. Sci. 105, 17029–17033 (2008).

23. Love, N. L. R. & Mazer, S. J. Region-specific phenological sensitivities and rates of climate warming generate divergent temporal shifts in flowering date across a species’ range. Am. J. Bot. 108, 1873–1888 (2021).

24. Park, D. S. et al. Herbarium specimens reveal substantial and unexpected variation in phenological sensitivity across the eastern United States. Phil. Trans. R. Soc. B 374, 20170394 (2018).

25. Preve y, J. et al. Greater temperature sensitivity of plant phenology at colder sites: implications for convergence across northern latitudes. Glob. Change Biol. 23, 2660–2671 (2017).

26. Richardson, B. A., Chaney, L., Shaw, N. L. & Still, S. M. Will phenotypic plasticity affecting flowering phenology keep pace with climate change? Glob. Change Biol. 23, 2499–2508 (2017).

27. Urban, M. C. Accelerating extinction risk from climate change. Science 348, 571–573 (2015).

28. Fordham, D. A. et al. How complex should models be? Comparing correlative and mechanistic range dynamics models. Glob. Change Biol. 24, 1357–1370, (2018).

29. Mazer, S. J. Ecological, Taxonomic, and Life History Correlates of Seed Mass Among Indiana Dune Angiosperms. Ecol. Monogr. 59, 153–175 (1989).

30. Bolmgren, K. & D. Cowan, P. Time – size tradeoffs: a phylogenetic comparative study of flowering time, plant height and seed mass in a north-temperate flora. Oikos 117, 424–429 (2008).

31. Jia, P., Bayaerta, T., Li, X. & Du, G. Relationships between flowering phenology and functional traits in eastern Tibet alpine meadow. Arct. Antarct. Alp. Res. 43, 585–592 (2011).

32. Metz, J. et al. Plant survival in relation to seed size along environmental gradients: a long-term study from semi-arid and Mediterranean annual plant communities. J. Ecol. 98, 697–704 (2010).

33. Molau, U. Relationships between flowering phenology and life history strategies in tundra plants. Arct. Antarct. Alp. Res. 25, 391–402 (1993).

34. Molau, U., Nordenha ll, U. & Eriksen, B. Onset of flowering and climate variability in an alpine landscape: a 10-year study from Swedish Lapland. Am. J. Bot. 92, 422–431 (2005).

35. Pangtey, Y. P. S., Rawal, R. S., Bankoti, N. S. & Samant, S. S. Phenology of high-altitude plants of Kumaun in Central Himalaya, India. Int. J. Biometeorol. 34, 122–127 (1990).

36. Cahill, A. E. et al. How does climate change cause extinction? Proc. R. Soc. B: Biol. Sci. 280, 20121890 (2013).

37. Thomas, C. D., Franco, A. M. A. & Hill, J. K. Range retractions and extinction in the face of climate warming. Trends Ecol. Evol. 21, 415–416 (2006).

38. Hargreaves, A. L. & Eckert, C. G. Evolution of dispersal and mating systems along geographic gradients: implications for shifting ranges. Funct. Ecol. 28, 5–21 (2014).

39. Brito-Morales, I. et al. Climate Velocity Can Inform Conservation in a Warming World. Trends Ecol. Evol. 33, 441–457 (2018).

40. Amano, T. et al. Links between plant species’ spatial and temporal responses to a warming climate. Proc. R. Soc. B: Biol. Sci. 281, 20133017 (2014).

41. Wiens, J. J. Climate-Related Local Extinctions Are Already Widespread among Plant and Animal Species. PLoS Biol. 14, e2001104 (2016).

42. Williams, J. W. & Jackson, S. T. Novel climates, no-analog communities, and ecological surprises. Front. Ecol. Environ. 5, 475–482 (2007).

43. Ackerly, D. D. Community Assembly, Niche Conservatism, and Adaptive Evolution in Changing Environments. Int. J. Plant Sci. 164, S165–S184 (2003).

44. Valladares, F. et al. The effects of phenotypic plasticity and local adaptation on forecasts of species range shifts under climate change. Ecol. Lett. 17, 1351–1364 (2014).

45. Park, D. S., Breckheimer, I. K., Ellison, A. M., Lyra, G. M. & Davis, C. C. Phenological displacement is uncommon among sympatric angiosperms. New Phytol. 233, 1466–1478 (2022).

46. Park, D. S. & Davis, C. C. Implications and alternatives of assigning climate data to geographical centroids. J. Biogeogr. 44, 2188–2198 (2017).

47. Li, D. et al. Climate, urbanization, and species traits interactively drive flowering duration. Glob. Change Biol. 27, 892–903 (2021).

48. Buri, A. et al. Soil factors improve predictions of plant species distribution in a mountain environment. Prog. Phys. Geogr. 41, 703–722 (2017).

49. Figueiredo, F. O. G. et al. Beyond climate control on species range: The importance of soil data to predict distribution of Amazonian plant species. J. Biogeogr. 45, 190–200 (2018).

50. Eyring, V. et al. Overview of the Coupled Model Intercomparison Project Phase 6 (CMIP6) experimental design and organization. Geosci. Model Dev. 9, 1937–1958 (2016).

51. Peters, G. P. et al. The challenge to keep global warming below 2 °C. *Nat*. Clim. Change. 3, 4–6 (2013).

52. Cai, L., Zhang, H. & Davis, C. C. PhyloHerb: A high-throughput phylogenomic pipeline for processing genome skimming data. Appl. Plant Sci. 10, e11475 (2022).

53. Marinho, L. C. et al. Plastomes resolve generic limits within tribe Clusieae (Clusiaceae) and reveal the new genus Arawakia. Mol. Phylogenet. Evol. 134, 142–151 (2019).

54. Weitemier, K. et al. Hyb-Seq: Combining target enrichment and genome skimming for plant phylogenomics. Appl. Plant Sci. 2, 1400042 (2014).

55. Park, B. & Donoghue, M. J. Phylogeography of a widespread eastern North American shrub, Viburnum lantanoides. Am. J. Bot. 106, 389–401 (2019).

56. Ruhfel, B. R., Gitzendanner, M. A., Soltis, P. S., Soltis, D. E. & Burleigh, J. G. From algae to angiosperms–inferring the phylogeny of green plants (Viridiplantae) from 360 plastid genomes. BMC Evol. Biol. 14, 23 (2014).

57. Kuznetsova, A., Brockhoff, P. B. & Christensen, R. H. B. lmerTest Package: Tests in Linear Mixed Effects Models. J. Stat. Softw. 82, 1–26 (2017).

58. Li, D. & Ives, A. R. The statistical need to include phylogeny in trait-based analyses of community composition. Methods Ecol. Evol. 8, 1192–1199 (2017).

59. Blomberg, S. P., Garland Jr, T. & Ives, A. R. Testing for phylogenetic signal in comparative data: behavioral traits are more labile. Evolution 57, 717–745 (2003).

60. Pagel, M. Inferring the historical patterns of biological evolution. Nature 401, 877–884 (1999).

61. Li, D., Dinnage, R., Nell, L. A., Helmus, M. R. & Ives, A. R. phyr: An r package for phylogenetic species-distribution modelling in ecological communities. Methods Ecol. Evol. 11, 1455–1463 (2020).

62. Nakagawa, S. & Schielzeth, H. A general and simple method for obtaining R^2^ from generalized linear mixed-effects models. Methods Ecol. Evol. 4, 133–142 (2013).

63. Robin, X. et al. pROC: an open-source package for R and S+ to analyze and compare ROC curves. BMC Bioinform. 12, 77 (2011).

64. Sing, T., Sander, O., Beerenwinkel, N. & Lengauer, T. ROCR: visualizing classifier performance in R. Bioinformatics 21, 3940–3941 (2005).

65. Anderson, J. T., Inouye, D. W., McKinney, A. M., Colautti, R. I. & Mitchell-Olds, T. Phenotypic plasticity and adaptive evolution contribute to advancing flowering phenology in response to climate change. Proc. R. Soc. B: Biol. Sci. 279, 3843–3852 (2012).

66. Franks, S. J., Sim, S. & Weis, A. E. Rapid evolution of flowering time by an annual plant in response to a climate fluctuation. Proc. Natl Acad. Sci. 104, 1278–1282 (2007).

67. Borges, R. M. Phenotypic plasticity and longevity in plants and animals: cause and effect? J. Biosci. 34, 605–611 (2009).

68. Baines, P. G. & Folland, C. K. Evidence for a rapid global climate shift across the late 1960s. J. Clim. 20, 2721–2744 (2007).

69. Turner, J., Overland, J. E. & Walsh, J. E. An Arctic and antarctic perspective on recent climate change. Int. J. Climatol. 27, 277–293 (2007).

70. Salmona, J., Heller, R., Quéméré, E. & Chikhi, L. Climate change and human colonization triggered habitat loss and fragmentation in Madagascar. Mol. Ecol. 26, 5203–5222 (2017).

71. Hesselbarth, M. H. K., Sciaini, M., With, K. A., Wiegand, K. & Nowosad, J. landscapemetrics: an open-source R tool to calculate landscape metrics. Ecography 42, 1648–1657 (2019).

